# A comparative single-cell transcriptomic atlas for diverse populations of vertebrate hair cells

**DOI:** 10.64898/2026.09.02.748811

**Authors:** Mahashweta Basu, Nesrine Benkafadar, Beatrice Milon, Carlo Colantuoni, Brian R. Herb, Amanda Ciani Berlingeri, Ivan A. Cruz, Joshua Orvis, Emilia Luca, Mitsuo P. Sato, Dunia Abdul-Aziz, Meghan Dewan, Kathleen Gwilliam, Gabriella Manilla, Christopher Shults, R. Shaun Adkins, Yang Song, Anup Mahurkar, John V. Brigande, Alain Dabdoub, Albert Edge, Andrew K. Groves, Ksenia Gnedeva, Stefan Heller, Ronna Hertzano, Tatjana Piotrowski, David W. Raible, Yehoash Raphael, Jennifer Stone, Litao Tao, Mark E. Warchol, Seth A. Ament, Lisa V. Goodrich

## Abstract

Mechanosensitive hair cells vary widely in morphology and regenerative capacity across vertebrate organs and species. To investigate their underlying transcriptomic diversity, we integrated human, mouse, chicken, and zebrafish single-cell and single-nucleus RNA sequencing datasets and assembled a cross-species atlas of hair cells spanning organs, developmental stages, and species. Analysis of 29 hair cell populations, encompassing the major cochlear, vestibular, and lateral-line hair cell types, identified approximately 5,000 genes enriched in at least one hair cell population compared to supporting cells from the same organs. Unsupervised clustering of these hair cell-enriched (HCE) genes defined species-, organ-, and hair cell state–associated cohorts as well as broadly conserved hair cell-enriched programs. Using an AUC-based scoring framework, we further defined 884 pan hair cell-enriched (pan-HCE) genes with elevated expression in most developing and/or mature hair cell populations, including genes implicated in deafness, mechanotransduction, and synaptic transmission, along with genes not previously linked to hair cell function. Independent analysis of developing hair cells using the same metrics stratified pan-HCE genes based on when they are first enriched and identified an additional 97 genes that are transiently enriched. We used the pan- and developing HCE gene sets to assess transcriptional similarity between baseline hair cell states and hair cells produced during avian hair cell regeneration and in mouse cochlear organoids, as well as hair cell-like populations produced by fibroblast reprogramming. HCE gene sets with different developmental dynamics identified young *vs.* more mature hair cells when projected onto independent single-cell RNA sequencing datasets from developing zebrafish, mouse, and human. We provide a web-based resource of all HCE metrics and expression profiles, enabling future exploration of vertebrate hair cell gene expression across organs, species, and experimental contexts.

## Introduction

Vertebrate hair cells convert mechanical stimuli into electrical signals and are essential for hearing and balance (Hudspeth, 2008)(Qiu and Müller, 2022)(Fettiplace and Hackney, 2006). Hair cells convert mechanical deflections of their stereocilia into electrical signals that are communicated through synapses with primary afferent neurons to the central nervous system (Ó Maoiléidigh and Ricci, 2019). Vertebrates have many types of hair cells, distributed across several sensory epithelia and defined by their unique morphology, position, and functional specializations (**Fig. 1A**). In mammals, hearing depends on inner and outer hair cells in the organ of Corti in the cochlea, while in birds, hearing relies on tall and short hair cells in the basilar papilla. Fish lack a dedicated hearing organ, but they detect sound with hair cells in the saccule (Schulz-Mirbach and Ladich, 2016)(Popper and Fay, 1993). In all species, hair cells in the vestibular sensory organs of the utricle, saccule, and ampullae (the maculae and cristae) detect information about head position and movement (Khan and Chang, 2013). Additionally, fish and amphibians sense fluid motion across their bodies using hair cells in the lateral line system, which are composed of sensory organs called neuromasts (Bleckmann and Zelick, 2009).

**Figure 1:**
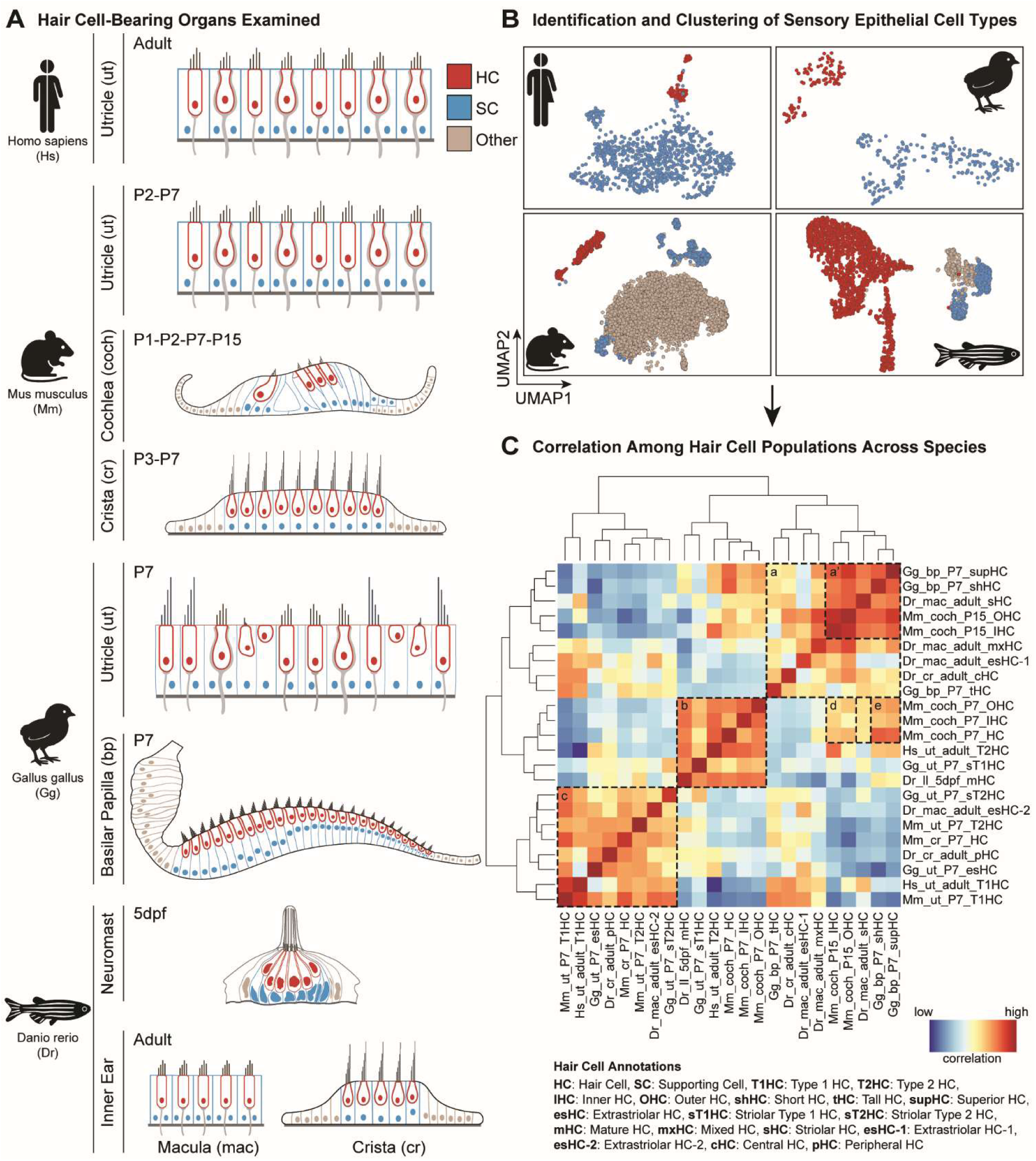
Diversity among sensory hair cells from different species, organs, and developmental stages. **(A)** Diagrammatic views of cross-sections through a variety of sensory epithelia that contain hair cells (red) and supporting cells (blue) in human (utricle), mouse (organ of Corti in the cochlea, crista, utricle), chicken (basilar papilla, utricle), and zebrafish (neuromast, macula, and crista). The age of the animal at the time of cell collection is indicated. (**B**) Examples of t-SNE and UMAP plots that were used to identify cell types from these tissues, with hair cells in red, supporting cells in blue, and other cell types in grey. (**C)** Heatmap showing correlations among the most mature hair cell types that were calculated by a MetaNeighbor analysis. Boxes a-e indicate notable correlation patterns, with a’ corresponding to a smaller branch within the larger “a” branch. The authors’ original annotations were used to name each population.

In all hair cell-containing epithelia, hair cells are surrounded by non-sensory supporting cells that are derived from a shared developmental lineage but exhibit distinct morphologies and functions relative to hair cells (Wan et al., 2013)(Guo et al., 2024). When hair cells die in non-mammalian vertebrates, supporting cells replace them either by mitotic regeneration or a non-mitotic process called direct transdifferentiation (Stone and Cotanche, 2007). Supporting cells can regenerate a fraction of one type of hair cells in vestibular epithelia of mammals but not in the cochlea (Kawamoto et al., 2009)(Golub et al., 2012).

Gaining a deeper knowledge of the genes that are enriched in mature hair cells would provide insights into the unique gene expression programs that distinguish them from supporting cells and endow them with functional specializations, such as their stereocilia bundles. A defined set of hair cell-enriched (HCE) genes would provide a benchmark to help investigators gauge the extent to which regenerated hair cells mature and to deduce trajectories by which supporting cells give rise to hair cells, which could reveal potential targets for therapeutic intervention. Similarly, a list of HCE genes would inform on the efficacy of reprogramming toward the hair cell fate in a variety of contexts, including human cell-derived inner ear organoids. Because studies on hair cell biology, development, and regeneration are conducted across several animal types, a list of genes enriched in hair cells across species is needed to enable direct comparisons and hence accelerate the identification of therapeutically relevant mechanisms.

Previous cross-species comparisons of hair cell transcriptomes revealed core gene sets but did not consider the relationship with supporting cells, which is key for uncovering relevant developmental pathways. Additionally, only a fraction of hair-cell epithelia and developmental stages have been compared, limiting the ability to define shared modules or cellular homologies, especially since stage-specific transcriptional differences may obscure important biological convergences. To generate a comprehensive atlas of a wide variety of hair cells from different ages, tissues, and species, we examined single cell transcriptomes of hair cells and supporting cells in the inner ear organs of zebrafish, chickens, mouse, and humans, and in the zebrafish lateral line, using published and unpublished datasets. Using a binary classifier represented by AUC scores, we found ∼5000 genes that are enriched in at least one kind of hair cell. Although hair cells tended to share more transcriptional similarity within species than across species, we defined a cohort of ∼1000 genes that are consistently enriched in mechanosensitive hair cells relative to age- and organ-matched supporting cells across different epithelia and species. Many genes in this “pan-HCE” gene set encode proteins that are enriched in the stereociliary bundle. Moreover, the list included 25% of all known human deafness genes. Assessment of independent datasets confirmed that this panel of pan-HCE genes show enriched expression in developing, regenerating, and mature hair cells across species, in cochlear organoid-derived hair cells, and in skin fibroblasts during reprogramming towards the hair cell fate. Additionally, developmental differences in the onset of expression were preserved across species. This gene set enables a rich assessment of hair cell identity and can serve as a reference for investigators of hair cell biology. We have made the entire dataset available to the community for customized searches through a ShinyApp and the Gene Expression Analysis Resource portal (gEAR)(umgear.org).

## Results

### A comprehensive catalog of hair cell-enriched genes across time points, organs, and species

To gain insight into transcriptional diversity among hair cells, we assembled published scRNA-seq datasets from various hair cell-containing organs across four species and eleven developmental timepoints (**Fig. 1A, Table S1**). A human study profiled the utricle of adults with vestibular schwannomas undergoing surgical tumor resection (Luca et al., 2025). Mouse datasets included the cochlea at postnatal day 1 (P1), P2, P7, and P15; the utricle at P2 and P7; and the crista at P3 and P7 (Kolla et al., 2020)(Kalra et al., 2020) (Ranum et al., 2019)(Wilkerson et al., 2021). Although mouse sensory organs are still immature at these postnatal stages, many hair cells are already capable of mechanotransduction (Lelli et al., 2009)(Géléoc and Holt, 2003). Chicken datasets profiled the hearing organ (basilar papilla) and the utricle at P7 (Janesick et al., 2021)(Scheibinger et al., 2022). Zebrafish datasets included the lateral line (neuromasts) and inner ear (maculae of the utricle and saccule, cristae) at five days post fertilization (dpf) and in adults (Baek et al., 2022)(Shi et al., 2023)(Sandler et al., 2025). Zebrafish utricle and saccule hair cells were combined into a single “maculae” group (Baek et al., 2022)(Shi et al., 2023). Hair cells in chicken and zebrafish organs are generally considered mature at these stages, although some degree of ongoing hair cell addition persists except in the chicken basilar papilla (Brignull et al., 2009). Most datasets were produced from dissociated whole cells using the 10x Genomics 3’ Gene Expression platform, while a few utilized SMART-seq full transcript scRNA-seq or single-nuclei RNA sequencing.

From these datasets, we extracted 43 populations of hair cells based on the authors’ original annotations (examples are provided in **Fig. 1B**). Populations corresponded to molecularly defined clusters in each dataset and were categorized by species, organ, age, and well-defined subtypes within each organ (e.g., inner versus outer hair cells in the mammalian cochlea) (**Table S1**). 23 populations were classified as being capable of mechanotransduction and approaching functional maturity; we defined these as “mature” though in some cases, additional maturation may occur at later ages. An additional 6 groups correspond to hair cells that are mechanosensitive but still maturing. The remaining 14 groups were classified as immature hair cells, as most or all lacked mechanical sensitivity. For initial analyses, we set aside the 20 immature and maturing subtypes to focus on genes associated with mature functions.

Possible cellular homologies among the 23 populations of mature hair cells were evaluated using MetaNeighbor (Crow et al., 2018), which calculates pairwise similarity across cell clusters and performs hierarchical clustering. This analysis supported many biologically meaningful relationships but did not identify any previously unknown relationships between known hair cell subtypes (**Fig. 1C**). Hair cell populations generally clustered by sensory modality. For instance, P15 mouse cochlear hair cells clustered with hair cells from the chicken basilar papilla (cluster a’), perhaps reflecting similar roles in audition. However, we found no support for finer homology of organ-specific types of hair cells across species, as neither of the specific mouse cochlear hair cell subtypes (inner or outer) showed greater homology with any of the chicken basilar papilla subtypes (tall, superior, or short). Hair cells from zebrafish maculae were grouped within a broader cluster that also included P15 mouse cochlea and chicken basilar papilla (cluster a). Apart from human type II hair cells, vestibular hair cells of the different species generally clustered together (cluster c), but not by organ identity. The low correlation between human type II hair cells and those from other species aligns with previous human-mouse cross-species comparisons that also reported divergence (Luca et al., 2025)(Wang et al., 2024a). Age drove some clustering, with the P7 mouse cochlea (before hearing onset) segregated from the P15 mouse cochlea (after hearing onset) (cluster a vs. cluster d). In fact, the correlation between mouse cochlea hair cell clusters across development (P7 vs. P15, cluster d) was relatively modest compared to that between hair cells in the P7 chicken basilar papilla and P7 mouse cochlea (cluster e). Overall, the MetaNeighbor detected patterns suggest that many of the genes distinguishing these 23 populations of hair cells involve factors related to age, species, and sensory modality.

To catalog similarities and differences in gene expression programs across diverse hair cells, we sought to identify genes associated with the 23 mature hair cell populations and the 6 additional populations capable of mechanotransduction but not yet mature; these 29 populations differ by species, organ, subtype, and developmental age. 14 developing hair cell populations not yet capable of mechanotransduction were set aside for later comparison. Achieving comparability across datasets posed several challenges beyond those encountered in typical differential gene expression analyses:

#### i. Choice of comparison cell group

While all datasets profiled hair cells, there was considerable variation in the other cell types present, both for biological reasons (heterogeneity in the cellular makeup of each organ) and for more technical factors (differences in dissection and enrichment strategies). To ensure consistency, we assessed hair cell enrichment relative to supporting cells, which are epithelial cells that either surround or are in close proximity to hair cells (**Fig. 1A**). Supporting cells were chosen because they were present in all datasets, are morphologically and physiologically distinct from hair cells, and share a developmental origin with hair cells. Supporting cell subtypes were combined into a single reference group within each dataset to avoid confounding differences among subtypes. Specifically, we sought to find genes generally enriched in hair cells compared to any nearby supporting cell, as opposed to one specific subtype of supporting cell.

#### ii. Cross-species gene mapping

Because comparisons spanned four species, we generated a unified set of gene orthologs. We linked all mouse, chicken, and zebrafish orthologs with each human gene, based on sequence similarity and chromosomal location. Multiple alignments of orthologs are common, especially in zebrafish, where ∼25% of the genes are duplicated relative to tetrapods. In these cases, each human gene was linked separately to each ortholog. We performed manual annotation of chicken orthologs for established hair cell genes to address annotation gaps in the chicken genome. This process yielded 45,050 orthologous gene relationships, corresponding to 20,289 human genes (**Table S2**), which were used for downstream comparisons across conditions.

#### iii. Cross-platform differential expression metric

We sought a differential expression statistic that is robust to technological differences between datasets. The area under the receiver operator curve (AUC) is a measure of gene enrichment that effectively distinguishes cell types even when there is uncertainty about absolute expression levels (Crow et al., 2018)(Pullin and McCarthy, 2024). We opted to use AUC as a measure of how well a gene’s expression classifies whether a given cell is a hair cell or a supporting cell. An AUC score of 1 indicates a gene is a perfect predictor of cell identity, whereas a score of 0.5 indicates that the gene has no predictive ability. However, the standard AUC score does not distinguish whether a gene’s expression is enriched or depleted in hair cells relative to supporting cells, and scores between 0 and 0.5 are uninformative. To address this, we set AUC scores below 0.5 to 0.5. For genes enriched in supporting cells relative to hair cells, defined as those with log_2_ fold change [logFC] < 0, we reassigned the AUC score as 1 – AUC. This transformation produces scores between 0 and 1, with scores above 0.5 indicating enrichment in hair cells and scores below 0.5 indicating enrichment in supporting cells (**Fig. 2B,C**). We calculated AUCs to identify hair cell-enriched (HCE) genes in each of the 29 hair cell populations.

**Figure 2:**
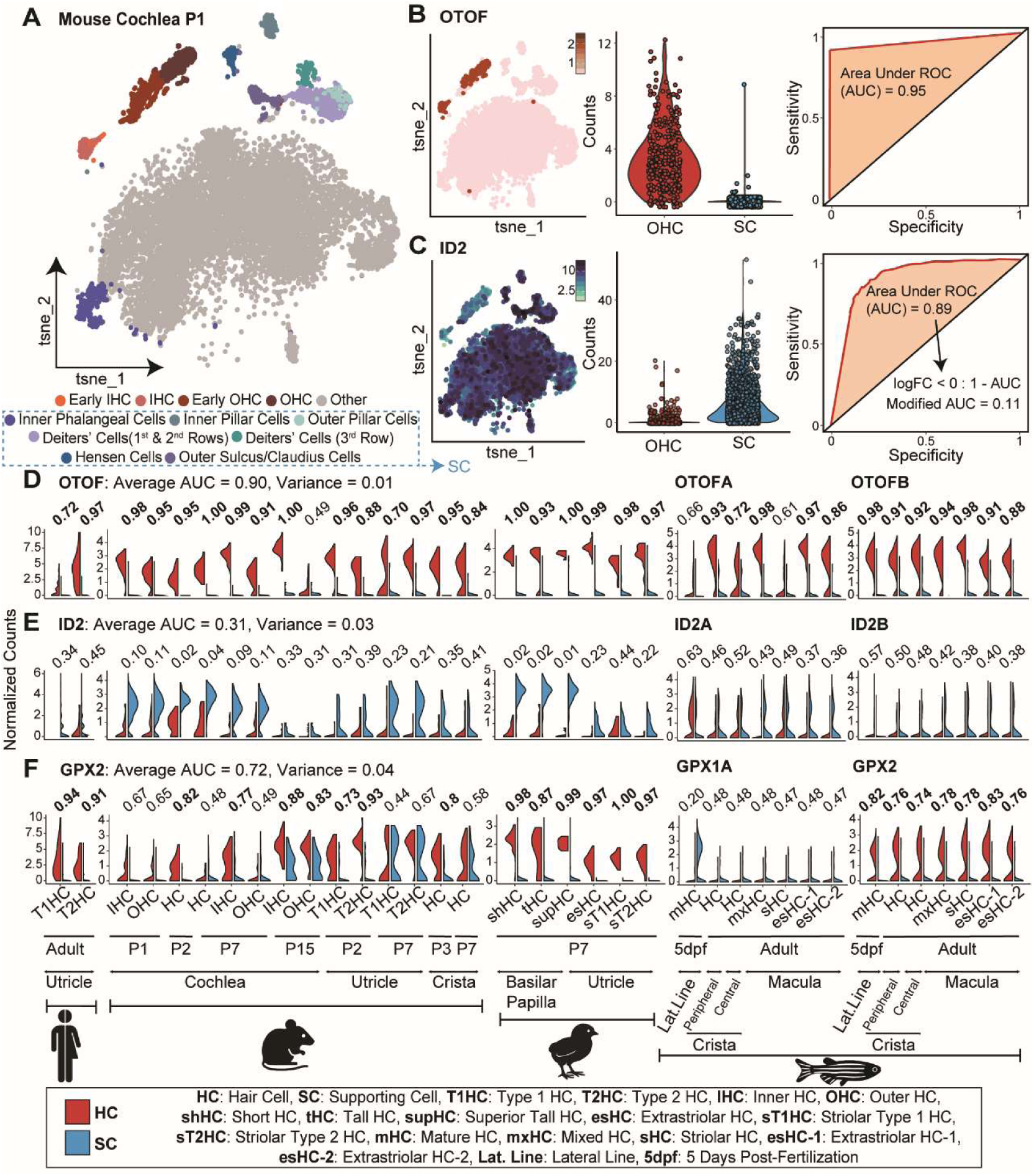
AUC values distinguish hair cells from supporting cells across datasets. (**A-C**) An example of how AUC values relate to gene expression levels, demonstrated using a dataset from the P1 mouse cochlea. As shown in a t-SNE plot, hair cells (red), supporting cells (blue), and other cell types (grey) were identified based on expression of known cell type markers such as *otoferlin* (*OTOF*) for hair cells (B) and *ID2* for supporting cells (C), shown both in feature plots (left) and violin plots (middle) of gene expression counts in outer hair cells (OHC) and in pooled supporting cells (SC), which consists of supporting cell types indicated by the dashed box in A. Based on these counts, a Receiver Operating Characteristic (ROC) curve (red) was plotted for each gene (right), with the Area Under ROC (AUC) marked in orange. For genes that are more highly expressed in SCs, the AUC was modified by subtracting from 1. This approach allowed us to focus on the hair cell/supporting cell relationship without any complications from gene expression in other cell types, such as what is observed for *ID2*. (**D-F**) Violin plots for *OTOF* (D), *ID2* (E), and *GPX2* (F) in all 29 hair cell populations, with corresponding AUC values above. Both zebrafish paralogs of each human gene are shown, so the zebrafish hair cell populations are duplicated. The name, age, organ, and species of each hair cell population, per the authors’ original annotations, is indicated below.

Genes with established unique functions in hair cells or supporting cells were used to validate the annotations of hair cell and supporting cell groups, as well as the reliability of AUC scores to assess differential gene expression. For example, in the neonatal mouse cochlea, *otoferlin* (*OTOF*), which is required for ribbon synapse function (Pangršič et al., 2012), was enriched in hair cells, while *inhibitor of differentiation 2* (*ID2*), which maintains the supporting cell fate during development (Jones et al., 2006), was enriched in supporting cells (**Fig. 2A-C**). Indeed, *OTOF* was enriched in most or all hair cell populations (**Fig. 2D**), whereas *ID2* was enriched in supporting cells from most datasets (**Fig. 2E**). Note that, by comparing only to supporting cells within the same dataset, our method avoids complications from gene expression in other cell types that might obscure important roles in hair cells. Thus, ID2’s AUC score is not influenced by its expression in other cochlear cell types (**Fig. 2C**). This approach is appropriate for our primary goal, which is to find genes that distinguish hair cells from supporting cells, even if those genes are expressed in other cells in the cochlea.

Although the absolute raw count values for transcripts differed widely across datasets, reflecting differences in technology, AUCs were consistently high for *OTOF*, with an average of 0.9 ± 0.01 across all 29 hair cell populations, while consistently low for *ID2*, with an average AUC of 0.31 ± 0.03. Thus, AUC is a reliable metric of hair cell enrichment, independent of technical factors. This approach also identified genes that showed enrichment only in a subset of hair cell groups, resulting in slightly lower average AUCs. For instance, many but not all hair cell populations had high expression of *glutathione peroxidase 2* (*GPX2*)(Chu et al., 1993), which encodes an antioxidant enzyme that contributes to oxidative stress responses, though no function in hair cells has been described (**Fig. 2F**). In this case, the average AUC was 0.72 ± 0.04, with values ranging from 0.2 to 1. To identify the genes most enriched in hair cells and also capture the expected diversity among hair cells, we defined hair cell-enriched genes in each dataset at a stringent AUC threshold ≥ 0.7. This analysis yielded 4,964 distinct genes that were strongly enriched in at least one of the 29 hair cell populations (**Table S2**). This catalog of HCE genes, which spans ages, organs and species, provides a useful resource for comparative studies of hair cell biology.

### Transcriptional variation across hair cells from different species, organs, and ages

To identify similarities and differences in gene expression programs across mechanically sensitive hair cells in an unbiased fashion, we performed hierarchical clustering of AUC scores. This approach computationally grouped genes with similar patterns of enrichment across all 29 groups of mechanically sensitive hair cells, resulting in 22 clusters of highly correlated genes (**Fig. 3A**, **Table S2, Figure 3–figure supplement 1**) that ranged in size from 12 human genes (clusters 16 and 21) to 1,831 genes (cluster 2).

**Figure 3:**
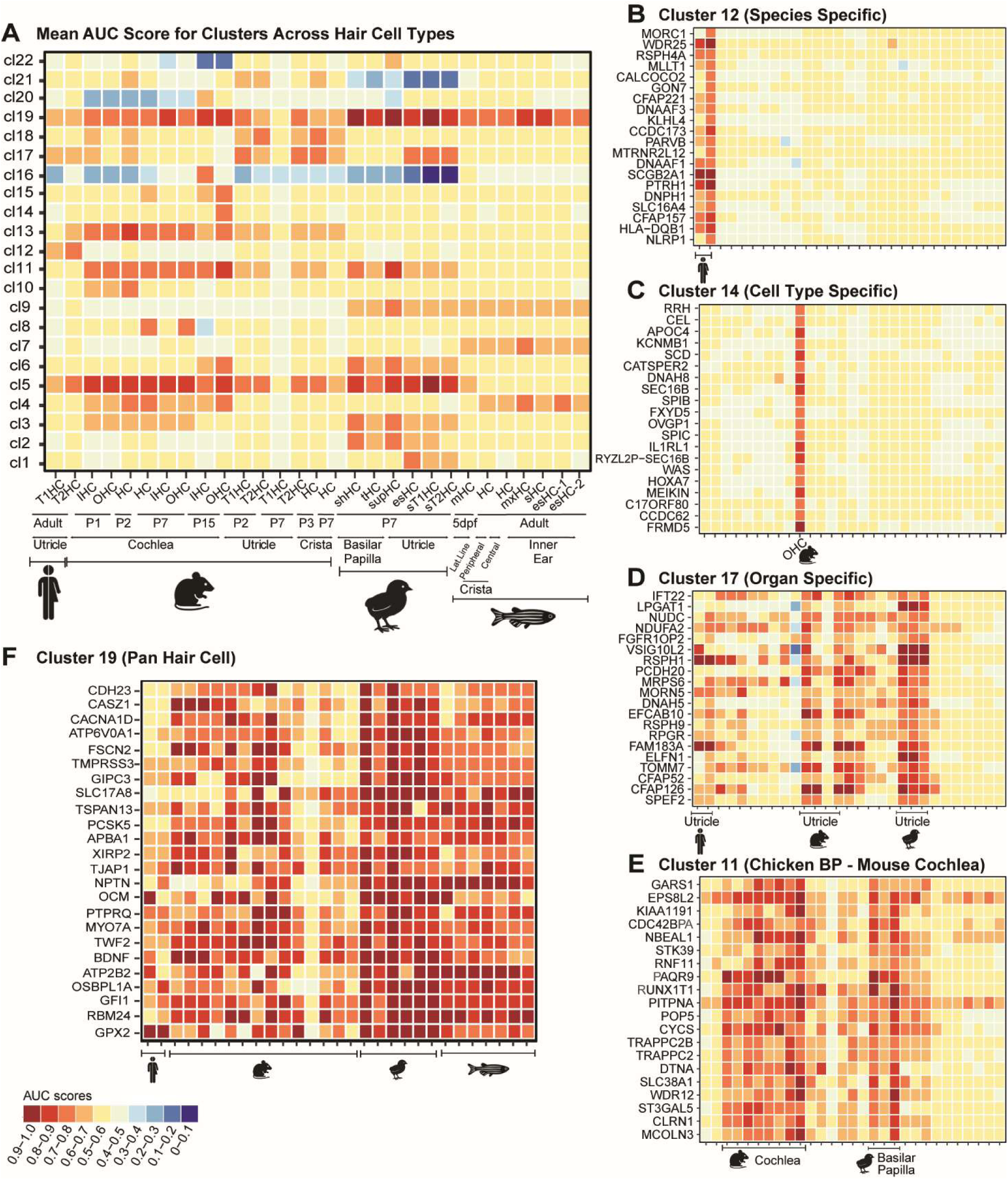
Hierarchical clustering reveals groups of hair cell-enriched genes with similar expression profiles across hair cell populations. **(A)** Heatmap showing the average AUC score for all genes in each of 22 clusters (cl1 to cl22) across the 29 hair cell populations, annotated below. **(B-F)** Heatmaps of AUC scores for the top 20 genes in clusters 12, 14, 17, 11, and 19 respectively. See Figure 3**–figure supplement 1** for additional information about clustering and Table S3 for information about the types of genes enriched in each cluster.

Examination of the clusters revealed several insights into how diverse hair cells differ from supporting cells and from each other. Since clusters are defined as sets of genes with similar patterns of enrichment relative to the other clusters, cluster identity was assessed by examining the average AUC value for all genes in that cluster in each of the 29 hair cell groups. We found that clusters differ substantially across species: 11 of the 22 clusters had higher AUC scores in a single species, namely clusters 1, 2, 7, 8, 10, 12, 14, 15, 16, 18, and 20. For instance, the average AUC value for genes in cluster 12 was higher in human utricle hair cells than in other hair cell groups (**Fig. 3B**). Other clusters showed high mean AUCs in hair cells from zebrafish (cluster 7) or chicken (clusters 1 and 2) (**Fig. 3A**). Among the mouse-dominated clusters, cluster 15 contained genes most enriched in cochlear hair cells, while clusters 14 and 16 were even more specific, corresponding to P15 mouse outer and inner hair cells, respectively (**Fig. 3A, C**). One of the genes in cluster 14 was SLC26A5 (prestin), which is essential for OHC function, underscoring the likelihood that other genes in that cluster are functionally important (J. Zheng et al., 2000)(Liberman et al., 2002)(Dallos et al., 2008). There was also a cluster driven by higher expression in mouse vestibular organs (cluster 18). Gene set enrichment analyses of the 22 co-expression clusters suggested that hair cells in different organs and species can be distinguished by quantitative differences in the expression levels of genes associated with energy metabolism and other homeostatic functions (**Table S3).** By contrast, genes related to core aspects of hair cell function, such as stereocilia function or calcium signaling, were mainly enriched in clusters of genes with high enrichment in many hair cell groups. These results suggest that aspects of hair cell function are defined by sets of genes with distinct expression in each species, but many genes involved in the core functions of hair cells are shared widely.

Eight clusters (clusters 3, 4, 9, 11, 13, 17, 21, and 22) were characterized by similar gene expression patterns across multiple (but not all) species. As expected, human utricular hair cells showed similarities to mouse cochlear and utricular hair cells (cluster 13, **Fig. 3A**). The analysis also revealed relationships between more evolutionarily divergent species, in some cases suggesting cross-species organ-specific enrichment. For instance, genes in cluster 17 were enriched in utricular hair cells from human, mouse, and chicken, highlighting a common set of genes related to vestibular function (**Fig. 3D**) and echoing relationships revealed by the MetaNeighbor comparisons (**Fig. 1C**). Many genes in this cluster are involved in the formation or motility of kinocilia, consistent with recent analysis of vestibular hair cells (Xu et al., 2026). On the other hand, cluster 11 consisted of 74 genes with higher AUC scores in organs associated with hearing, namely mouse cochlear and chicken basilar papilla hair cells, with slightly less enrichment in vestibular hair cells from both species (**Fig. 3E**). There were also clusters containing hair cells from mouse and zebrafish (cluster 4); chicken and zebrafish (cluster 9); and human, mouse, and zebrafish (cluster 21). Cluster 22 was characterized by lower AUC values across the board, with possible supporting cell enrichment in some mouse populations. The underlying associations amongst cells in each of these clusters were unclear.

Finally, three clusters stood out for expression in most or all hair cells in the analysis (clusters 5, 6, and 19). These clusters, which contained a total of 273 genes, differed from each other modestly. For instance, genes in cluster 5 tended to be less enriched in zebrafish, whereas cluster 19 stood out for consistently high mean AUCs across all 29 hair cell populations (**Fig. 3A, F**). Cluster 6 showed moderate enrichment across populations with particularly high AUC values in mature mouse outer hair cells and chicken short hair cells and striolar hair cells, suggesting possible shared features among these specific types. The higher AUC values in P15 mouse hair cells compared to earlier stages suggests this cluster might reflect functional maturity, for instance compared to cluster 5. Many of the genes in these three clusters are important for hair cell mechanotransduction, such as *CDH23* (encoding the tip-link protein Cadherin-23, cluster 19) (Bork et al., 2001)(Bolz et al., 2001), *TMC1* (encoding the mechanotransduction channel component Transmembrane Channel-like 1, cluster 6) (Pan et al., 2018), and *ESPN* (encoding the actin-bundling protein Espin, cluster 5) (L. Zheng et al., 2000), providing confidence that our approach can identify cohorts of genes associated with shared functional features that distinguish hair cells from supporting cells across species.

### Identification of genes enriched in most or all hair cells

Building on the identification of genes consistently enriched in hair cells vs. supporting cells by hierarchical clustering, we used a machine learning strategy to define a set of “pan-hair cell” genes that are enriched in most or all hair cell subtypes (**Fig. 4A**; details in Methods). Our goal was not to identify all hair cell genes or to find genes that distinguish hair cells from all other cell types, but instead to produce a list that reliably distinguishes hair cells from supporting cells across experimental contexts. Towards this end, we summed the AUC scores across groups to produce an aggregate score for each gene, leading to a ranking from the most to least hair-cell enriched. We compared these hair cell enrichment rankings to lists of reference genes shown to be highly expressed in hair cells relative to supporting cells in prior studies (i.e., “gold standards” according to standard machine learning terminology), including lists derived from independent bulk RNA-seq, scRNA-seq, and proteomics experiments (Scheibinger et al., 2022) (Tao et al., 2021)(Liu et al., 2018)(Krey and Barr-Gillespie, 2019)(Wilmarth et al., 2015)(Sprague et al., 2003), as well as a manually-curated gene list based on our field’s research on hair cell development and regeneration (**Table S4**). We used precision-recall statistics to set a threshold that balances the inclusion of as many reference genes as possible while minimizing false positives. To ensure robustness, we applied four alternative strategies to calculate the aggregate enrichment scores (**Fig. 4A**). We either included all 29 hair cell groups capable of mechanotransduction (“mechano”) or limited the analysis to the 23 most mature hair cells (“mature”). Then, we computed either an unweighted average of the AUCs across all hair cell groups (Enrichment Score 1, “ES1”) or an adjusted average so that each species has equal weight (“ES2”), thereby accounting for the fact that the mouse is represented the most. Each approach led to slightly different rankings and thresholds, so we used the intersect of these gene lists to generate the final list. At the precision-recall break-even point, this analysis identified 884 consistently hair cell-enriched genes or “pan-HCE” genes (**Fig. 4B, Figure 4-figure supplement 2, Table S2**). Ranking these pan-HCE genes by their mechano-ES2 score (**Fig. 4C**; see **Figure 4 - figure supplement 1** for ranking using other scores) identified those genes with the highest and most consistent enrichment across hair cell populations.

**Figure 4:**
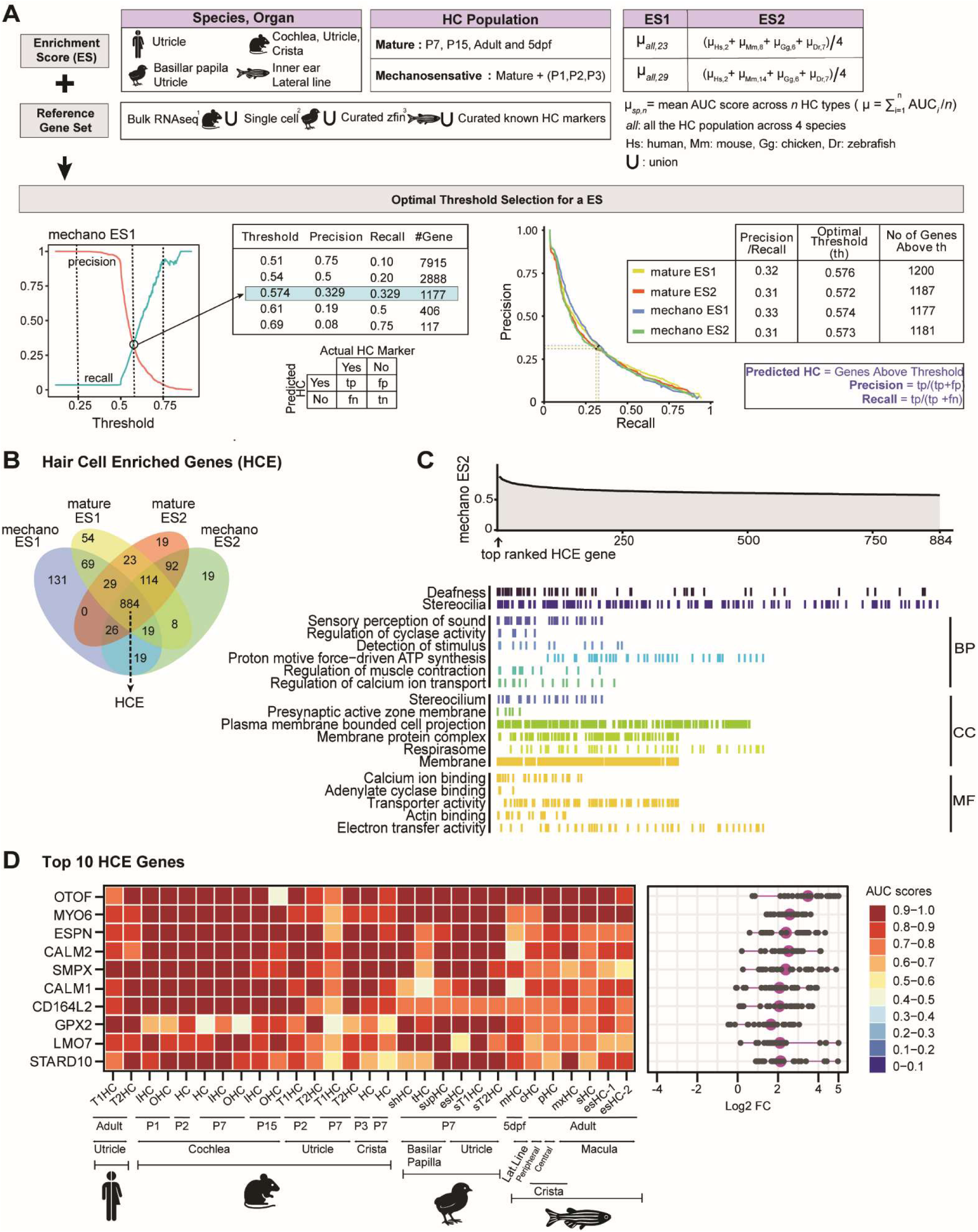
Identification of HCE genes using machine learning. **(A)** Schematic illustrating how four different scores were calculated from individual AUCs in diverse hair cell populations, which either included the most mature hair cells (“mature”) or all hair cells capable of mechanotransduction (“mechano”). An enrichment score (ES) was calculated either by averaging AUCs in all mature (mature-ES1) or mechanosensitive (mechano-ES1) hair cells, or by weighting average AUC scores evenly across species (ES2) for each population of hair cells (“mature-ES2” and “mechano-ES2”). Machine learning was used to compare the scores for every gene in the genome against a reference gene set, which consisted of genes known to be hair cell-enriched based either on bulk RNA-sequencing, independent single cell RNA sequencing results, or curated lists. Machine learning identified the threshold at which precision equals recall for each set of scores, which behaved very similarly and identified ∼1200 genes over the threshold in each case. **(B)** Venn diagram showing how the sets of genes identified using each scoring method overlap with each other, including the 884 genes that were present in all sets. **(C)** A ranked plot of mechano-ES2 scores for these 884 genes, with annotations indicating the positions of genes associated with deafness, stereocilia, and the top enriched GO terms. See Figure 4**–figure supplement 1** for rankings by the other three scores. (**D**) Heatmap displaying AUC scores for the top 10 HCE genes across 29 hair cell populations, accompanied by a dot plot showing their log fold-change values. See also Figure 4**– figure supplement 2** for a heatmap of all 884 AUC scores.

Manual examination of the pan-HCE genes supported the validity of our approach. The top ranked genes generally showed high enrichment across hair cells from different species, organs, and ages (**Fig. 4D, Figure 4--figure supplement 2, Table S2**). For instance, *XIRP2*, which encodes an actin binding protein localized to stereocilia (Scheffer et al., 2015), had AUC scores ranging from 0.57 (in mouse P7 utricle Type I hair cells) to 0.99 (in chicken P7 utricle striolar hair cells). Many genes with known roles in hair cells were among the top 50 pan-HCE genes, including genes that encode the hair cell-specific transcription factors POU4F3 (Erkman et al., 1996)(Xiang et al., 1997) and GFI1 (Wallis et al., 2003); hair cell-enriched unconventional myosins MYO7A, MYO6, and MYO15A (*54*); and OTOF (Pangršič et al., 2012). Structural components of stereocilia were strongly over-represented (odds ratio = 7.8, P = 3.8e- 104): the full list of 884 pan-HCE genes included 243 of 1,146 genes that encode for proteins localized to stereocilia (Krey et al., 2015), including *XIRP2*, *TWF2*, *PTPRQ*, and *FSCN2* among the top 50. Known genes linked to deafness (Walls WD, Azaiez H, Smith RJH., n.d.) were also enriched (odds ratio = 7.9, P = 1.0e-24), with 50 of 197 known human deafness genes annotated as pan-HCE genes, including *TMPRSS3*, *LHFPL5*, and *CIB2* among the top 50. As expected, pan-HCE genes included most or all genes from clusters 19 (**Fig. 3F**), 5, and other clusters enriched across hair cell groups (**Table S5**). This included cluster 11, which stood out for enhanced enrichment in mouse and chicken auditory organs, with notably lower AUC values in zebrafish (**Fig. 3E**). Of note, zebrafish AUC values were often lower, especially in those pan-HCE genes assigned to cluster 5. This is not unexpected, as a few HCE genes displayed lower AUC scores in some hair cell populations; we also refer to these as “pan-HCE” genes to reflect their consistent enrichment in the vast majority of hair cells across multiple hearing and balance organs from different species. Similarly, only 44% of the genes in cluster 6 were independently defined as HCE genes, likely due to the fact that that cluster is driven by enrichment in the more mature hair cell populations.

To gain additional insight into the types of genes that robustly distinguish hair cells from supporting cells, we used the Gene Set Enrichment Analysis (GSEA) algorithm to assess the pan-HCE gene list (**Fig. 4C**), weighting genes by their rank-order. GO terms for inner ear biological processes (e.g. “sensory perception of sound”, “detection of mechanical stimulus involved in sensory perception of sound”, “auditory receptor cell differentiation” and “equilibrioception”) and inner ear cellular components (e.g. “stereocilium”, “stereocilium tip”, “stereocilium bundle”) were significantly represented (**Fig. 4C, Table S6**) and those genes were among the most highly ranked on our list, as ordered by mechano-ES2 scores. Consistent with the role of the kinocilium in establishing and maintaining the hair bundle and the presence of actin-rich stereocilia in hair cells (and not supporting cells), GO terms associated with cilia and actin were also significantly represented, as were GO terms typically associated with neuronal functions, such as ion channels and synaptic vesicle proteins. This is unsurprising, as hair cells are secondary receptor cells that form synapses with afferent terminals from auditory, vestibular or lateral line neurons. By contrast, supporting cells are not capable of synaptic transmission. Finally, many pan-HCE genes were associated with GO terms related to energy metabolism (“respirasome”, “electron transfer activity”), likely reflecting the increased energetic requirements of hair cells compared to supporting cells.

### In situ hybridization validation of pan-hair cell gene expression

To examine how well the machine-learning analysis defined gene enrichment in hair cells relative to supporting cells across species and organs, we assessed expression of predicted pan-HCE genes using hybridization chain reaction (Choi et al., 2018), RNAscope (Benkafadar et al., 2024), or spatial transcriptomics in mouse cochlea, utricle, and crista; zebrafish lateral line neuromast, utricle, and crista; and chicken basilar papilla and utricle. 10 genes were selected for analysis based on their reliably high AUC scores in all species (**Figure 5-figure supplement 1**), their predicted role in the hair bundle or the synapse, and their predicted detectability based on expression levels in published and unpublished datasets. The analysis included six genes in cluster 5, even though this cluster was characterized by lower enrichment in zebrafish. For genes with two zebrafish paralogs, both zebrafish genes were assayed. One selected gene, *Tmem255b*, had no identifiable ortholog in zebrafish, so it was analyzed only in birds and mice.

**Figure 5:**
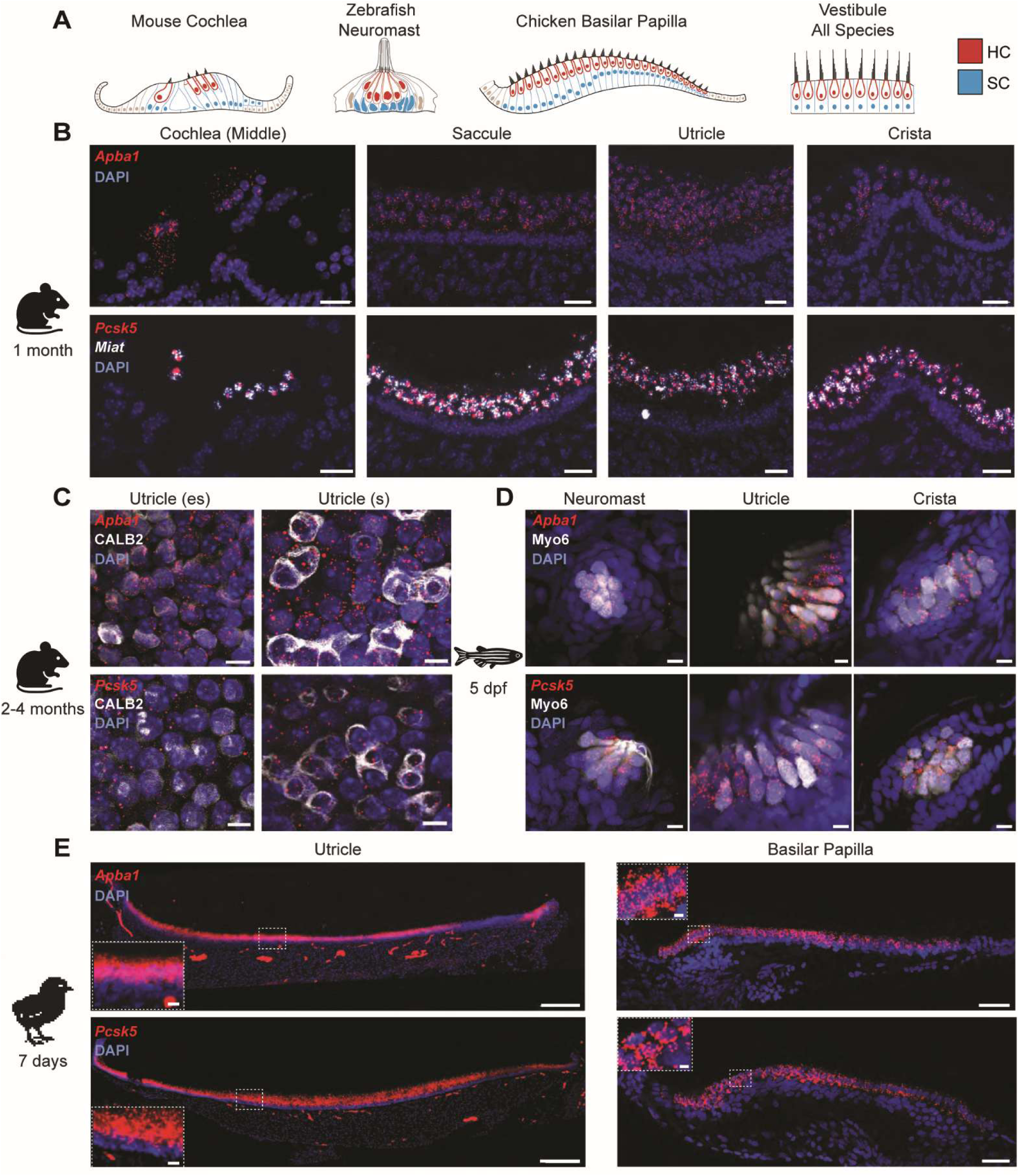
Pan hair-cell enrichment of *APBA1* and *PCSK5* across organs and species. (**A**) Illustrations of the sensory epithelia assayed by *in situ* hybridization. (**B**) RNAscope *in situ* hybridization confirming expression of *Apba1* (top) and *Pcsk5* (bottom) in sections through the adult mouse cochlea, saccule, utricle, and crista, as illustrated in A. *Miat* (white) marks hair cells. Scale bar: 20 µm. (**C**) Top down views of the extrastriolar (es) and striolar (s) regions of whole adult mouse utricles processed for HCR *in situ* hybridization of *Apba1* and *Pcsk5*. Calb2 immunostaining indicates Type I and Type II hair cells and the calyceal endings onto Type I hair cells. Scale bar: 6 µm. (**D**) HCR *in situ* hybridization of *Apba1* and *Pcsk5* in sensory organs of the lateral line (neuromast) and inner ear (utricle, crista) of 5 dpf zebrafish larvae, with hair cells independently marked by Myo6 immunostaining (white). Organs are oriented essentially as in A. Scale bar: 5 µm. **(E**) HCR *in situ* hybridization of the P7 chicken utricle and basilar papilla. Insets are high power views of the boxed regions, which illustrate enrichment of *Apba1* and *Pcsk5* (red) in hair cells and not in the underlying supporting cells, marked by DAPI. Scale bar: 100 µm (utricle), 50 µm (basilar papilla), 10 µm (utricle inset), and 20 µm (basilar papilla inset). See Figure 5**–figure supplement 1** for violin plots of all genes tested by *in situ* hybridization and Figure 5**–figure supplement 2** for spatial transcriptomic validation.

Seven of the ten genes selected for validation were detected consistently across tissues, with mRNA present in the hair cell cytoplasm and/or nucleus (**Fig. 5**). Only three genes (*Rab3ip, Runx1t1, Pgm2l1*) failed to show clear enrichment in hair cells relative to supporting cells in all organs in all three species. The remaining genes (*Apba1, Syt14/14b, Snap91, Pcsk5/5a, Fscn2/2a, Xirp2/2b, Tmem255b*) had consistently enriched hybridization signal in hair cells relative to supporting cells, shown for *Apba1* (also known as *Mint1*), which encodes a presynaptic component (Okamoto and Südhof, 1997), and *Pcsk5*, which encodes a member of the subtilisin-like proprotein convertase family and has been previously localized to zebrafish hair cells (Chitramuthu et al., 2010) (**Fig. 5B-E**). Consistent with the more neuronal nature of the hair cell as compared to a supporting cell, several genes were also enriched in spiral ganglion neurons, as determined both by RNAscope and using the Visium HD platform to perform spatial transcriptomics on adult mouse cochlea (**Figure 5–figure supplement 2**). Thus, our analysis uncovered broadly expressed hair cell genes that may encode proteins important for common hair cell functions such as mechanotransduction and synaptic transmission, and that can be used across species to distinguish hair cells from supporting cells.

### Expression patterns of pan-hair cell-enriched genes during development and regeneration

Given the emerging importance of cross-species comparisons in biology, it is valuable to have a large set of common markers whose expression captures the core features of a hair cell. This is particularly relevant to the field of hair cell regeneration, where expression of one or two markers such as Myo6 is insufficient to identify a cell as a “hair cell” following engineering of stem cells or expression of reprogramming factors. While species-specific markers often suffice, a large set of common genes enables comparison across species, especially if these genes are expressed at analogous stages of hair cell development and differentiation. To see if our list might provide a useful metric for the field, we examined pan-HCE gene expression in independent transcriptomic datasets that covered multiple species, developmental stages, and experimental conditions.

We first compared AUC scores for individual genes in E16 mouse cochlea, in which hair cells have begun to differentiate but are not yet mechanosensitive (Lelli et al., 2009), to the aggregate mechano-ES2 score for that gene in the cross-species comparison. We defined enrichment as an AUC>0.6, as this is just over the threshold defined by machine learning for all four scoring methods (**Fig. 4A**). This analysis showed that many pan-HCE genes (202/884) were enriched in hair cells by E16 (**Fig. 6A**, Quadrant (Q)1). In addition, we identified a set of genes with high AUC values at E16 that were not on the pan-HCE list (**Fig. 6A**, Q4), including *Atoh1*, which encodes a transcription factor essential for hair cell specification and differentiation (Cai and Groves, 2015). Thus, fewer pan-HCE genes are expressed developmentally than after the onset of mechanotransduction, raising the possibility that pan-HCE genes might also be useful for assessing the degree of hair cell maturity.

**Figure 6:**
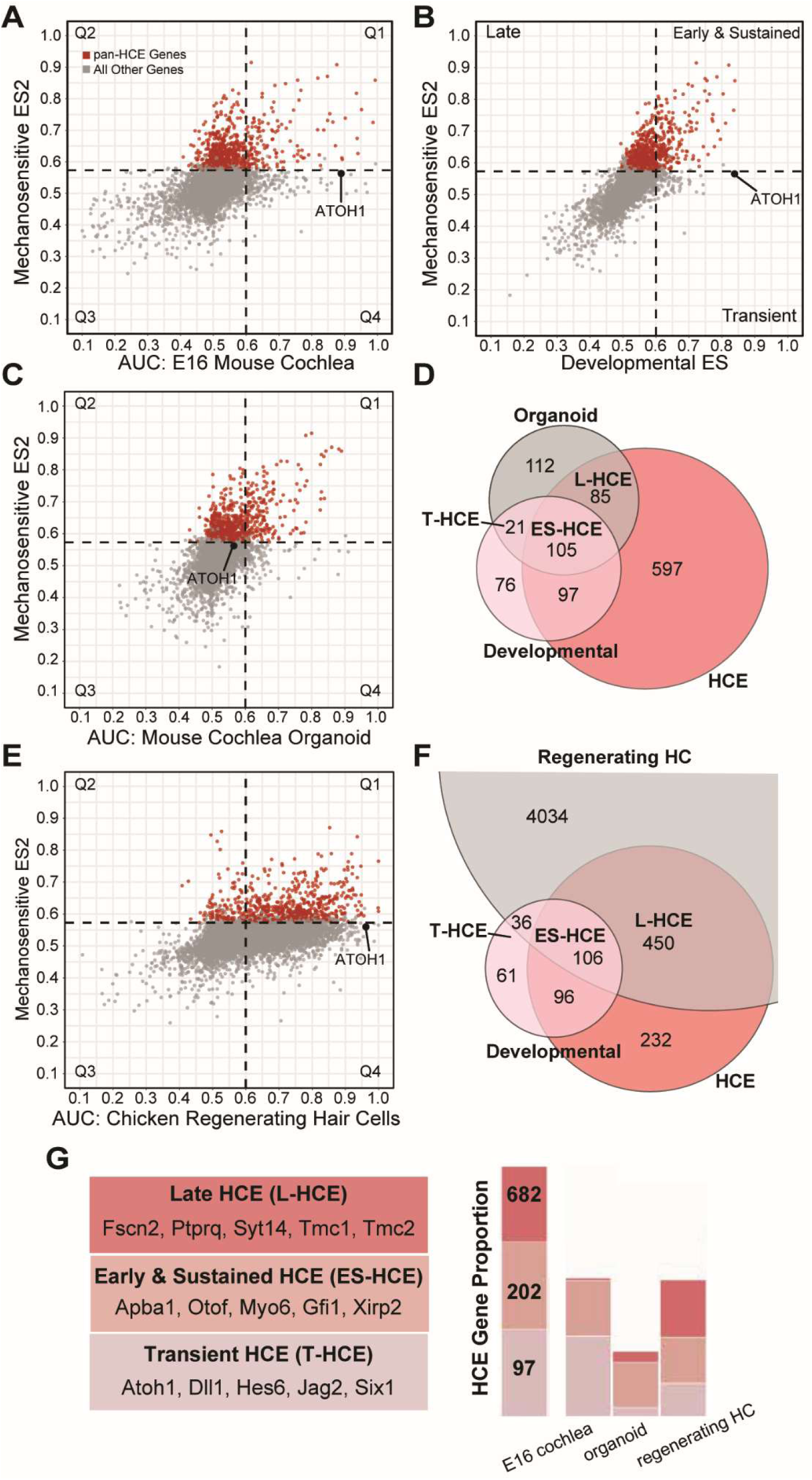
The use of HCE genes to assess hair cell development and regeneration *in vitro* and *in vivo*. (**A-C, E**) Scatter plots showing relationships between the mechano-ES2 score (y-axis) and: the mean AUC for hair cells in the E16 mouse cochlea (A), in 14 developing hair cell populations that were each compared to supporting cells in the same dataset (“Developmental ES”) (B), for 2 populations of hair cells from mouse cochlea organoids (C), and for newly regenerated hair cells in the chicken basilar papilla (E). Quadrants 1-4 are labeled in A (Q1-Q4) and B (late, early and sustained, transient, no label). Dashed lines indicate an AUC threshold of 0.6, which is close to the threshold determined by machine learning (Fig. 4). (**D, F**) Venn diagrams illustrating the overlap between genes enriched in organoid hair cells (D, gray) or regenerating chicken hair cells (F, gray) with the 884 HCE genes (“HCE”, red) and the enriched gene sets from developing hair cells (“Developmental”, pink). (**G**) Examples of genes classified as late, early and sustained, or transient and their proportions in hair cells from three sources. The complete HCE list includes 884 pan HCE genes and 97 transient HCE genes. The first stacked bar graph shows the total number of HCE genes in each category (late, early, transient). The remaining bars represent the proportion of genes in each of these categories that is significantly enriched in E16 mouse cochlea, cochlear organoid, and chicken regenerating hair cells. See also Figure 6**–figure supplement 1** for additional plots and Venn diagrams; **Table S9** for genes in each HCE subset; and **Table S10** for lists of genes at each intersection.

We reasoned that calculating enrichment scores for genes at early developmental stages could provide a broader assessment of hair cell identity. We therefore performed an independent analysis of gene enrichment in 14 immature hair cell types from all four species compared to supporting cells, calculating a “developmental enrichment score (ES)” for each gene as its average AUC across all 14 groups. Comparing mechano-ES2 to developmental ES scores stratified the original pan-HCE genes and added a third tier of genes with high AUCs only in developing hair cells (**Fig. 6B; Table S7, S8, S9**). Among the 884 pan-HCE genes (**Fig. 6B**, red), 202 genes had both high mechano-ES2 and developmental ES scores, suggesting they are enriched early and then sustained through maturity (**Fig. 6B, G**, early and sustained HCE, “ES-HCE”). The remaining 682 pan-HCE genes, on the other hand, had high mechano-ES2 and low developmental ES scores, indicating greater enrichment after the onset of mechanotransduction (**Fig. 6B, G**, late HCE, “L-HCE”). Of note, many L-HCE genes (235/682, 34.5%) were independently assigned to Cluster 6, lending support to the idea that Cluster 6 consists of genes needed for functional maturity. This analysis also revealed 97 genes with developmental ES scores greater than 0.6 and mechano-ES2 scores less than 0.6. These “transient HCE” (T-HCE) genes represent a younger hair cell state that is apparent across species, epitomized by *ATOH1* (**Fig. 6B, G**, transient HCE, “T-HCE”). In summary, this comparison identified a total of 981 HCE genes that can be used to identify hair cells across developmental stages and species, with 884 pan-HCE genes (202 ES-HCE + 682 L-HCE) and 97 T-HCE genes.

To evaluate whether known developmental differences can be detected by assessing HCE gene expression, we re-analyzed gene expression in organoids derived from postnatal LGR5+ supporting cells (McLean et al., 2017). Previous transcriptomic analysis identified two hair cell clusters (Kalra et al., 2023), with one being slightly more mature (cluster 8) than the other (cluster 7). We compared each hair cell cluster to the supporting cell clusters, as originally annotated (**Table S8)**. The resulting mean AUC scores for all organoid-derived hair cells were plotted against mechano-ES2 scores from the cross-species comparison (**Fig. 6C,D**) or developmental ES scores (**Figure 6–figure supplement 1**). Genes with AUCs >0.6 were considered to be enriched, as this is just above the thresholds identified by machine learning. This approach revealed 21/97 T-HCE genes (21.6%) and 105/202 ES-HCE genes (52.0%) that were also enriched in the organoids. By contrast, among the 682 L-HCE genes, only 85 (12.5%) had AUC scores >0.6 (**Fig. 6D**). Based on this analysis, hair cells generated in cochlear organoids more closely resemble young hair cells than mature hair cells. Consistent with the idea that this method can detect developmental differences, cluster 8 hair cells expressed fewer T-HCE genes (18.6% vs. 27.8%) and more ES-HCE + L- HCE genes (34.6% vs. 22.8%) than cluster 7 hair cells, which were independently determined to be less mature (**Figure 6–figure supplement 1**). Thus, HCE gene expression recapitulates developmental differences between cluster 7 and cluster 8 hair cells, confirming that our method reproduces the authors’ original conclusion (McLean et al., 2017) and thus validating its value as a metric. This analysis suggests that HCE gene sets can be used to compare methods of generating organoids and to refine culture conditions that optimize hair cell maturity.

To evaluate regenerated hair cells *in vivo*, we used the same approach on the chicken basilar papilla, where supporting cells produce new hair cells that gradually mature and eventually restore function over several weeks (Sato et al., 2024a). Transcriptomic analysis previously revealed that a population of newly regenerated hair cells is not yet equivalent to mature basilar papilla hair cells (Janesick et al., 2022), consistent with work from others (Stone et al., 1996). We therefore examined how this intermediate stage is reflected by HCE gene expression (**Fig. 6E,F, Figure 6–figure supplement 1, Table S7, S10**). Consistent with the observation that new hair cells are still maturing, 36 out of 97 T-HCE genes (37.1%), 106 out of 202 ES-HCE genes (52.5%), and 450 out of 682 L-HCE genes (66%) were highly enriched (**Fig. 6F**). Notably, HCE genes represented a small proportion of the genes with high AUC values in the newly regenerated chicken hair cells (Q1+Q4, **Fig. 6E**), emphasizing the fact that differences among species can obscure common gene cohorts.

The ability of supporting cells to transdifferentiate to hair cells in some species and organs has inspired great interest in identifying transcription factors that promote hair cell gene expression and might therefore reprogram supporting cells *in vivo*. Previous research identified a combination of four transcription factors that are sufficient to drive expression of large cohorts of hair cell genes in cultured fibroblasts (Menendez et al., 2020), as assessed by bulk RNA-seq of untransfected *vs.* transfected fibroblasts. Accordingly, we could not calculate AUCs for this dataset. Instead, we quantified the fold-change of gene expression between normal fibroblasts and fibroblasts that were reprogrammed by expression of *Gfi1*, *Atoh1*, *Pou4f*3, and *Six1* (**Table S7, S10**). We then compared the log_2_ fold change to the mechano-ES2 AUC for all genes in the genome. We found that many pan-HCE genes (254/884, 28.7%) were induced in the reprogrammed fibroblasts, and that this included few T-HCE genes (9/97, 9.3%), suggesting some degree of maturation. Likewise, fewer genes with high developmental ES were significantly induced (**Figure 6–figure supplement 1**). These findings suggest that HCE gene expression can be used to assess data for which there are no AUCs and that such an analysis can generate new hypotheses, such as the possibility that additional treatments may be needed to stimulate certain cohorts of genes associated with hair cell identity in fibroblasts.

To further assess the value of HCE gene sets for characterizing hair cell development and regeneration when AUC values are not readily available, we projected the T-HCE and L-HCE gene expression modules onto independent published datasets from zebrafish, mouse, and human that each contained multiple developmental or regenerating stages (**Fig. 7**)(Baek et al., 2022)(Sur et al., 2023)(Wang et al., 2024b)(van der Valk et al., 2023). Supporting cells and hair cells were identified using the authors’ annotations, including information about when the cells were profiled. As expected, UMAPs revealed clear trajectories from supporting cells to hair cells in the regenerating lateral line (**Fig. 7A-E**), from progenitors to hair cells in the developing zebrafish lateral line and otic vesicle (**Fig. 7F-J**) and in the developing mouse utricle (**Fig. 7K-M**). Across these diverse tissues, the T-HCE gene set was enriched in progenitors and immature hair cells and then downregulated in the more mature hair cells (**Fig. 7B, G, L**), which instead showed enriched expression of the L-HCE gene set (**Fig. 7C, H, M**). These patterns were clear not only in UMAPs, where the developmental progression is spatially organized, but also in violin plots, which illustrate the overall level of expression for each gene set in defined cell types (**Fig. 7D, E, I, J**). This approach revealed additional dynamics in expression in the regenerating lateral line, where the T-HCE gene set was relatively stable but the L-HCE gene set rapidly lost expression in mature hair cells and recovered by 5 hours after neomycin treatment (**Fig. 7D,E**). Similar dynamic changes in T-HCE and L- HCE gene set expression were observed in hair cells from fetal and adult human cochleae: T-HCE genes were high at fetal ages and low in the adult, whereas L-HCE genes were progressively enriched across development (**Fig. 7N-P**).

**Figure 7:**
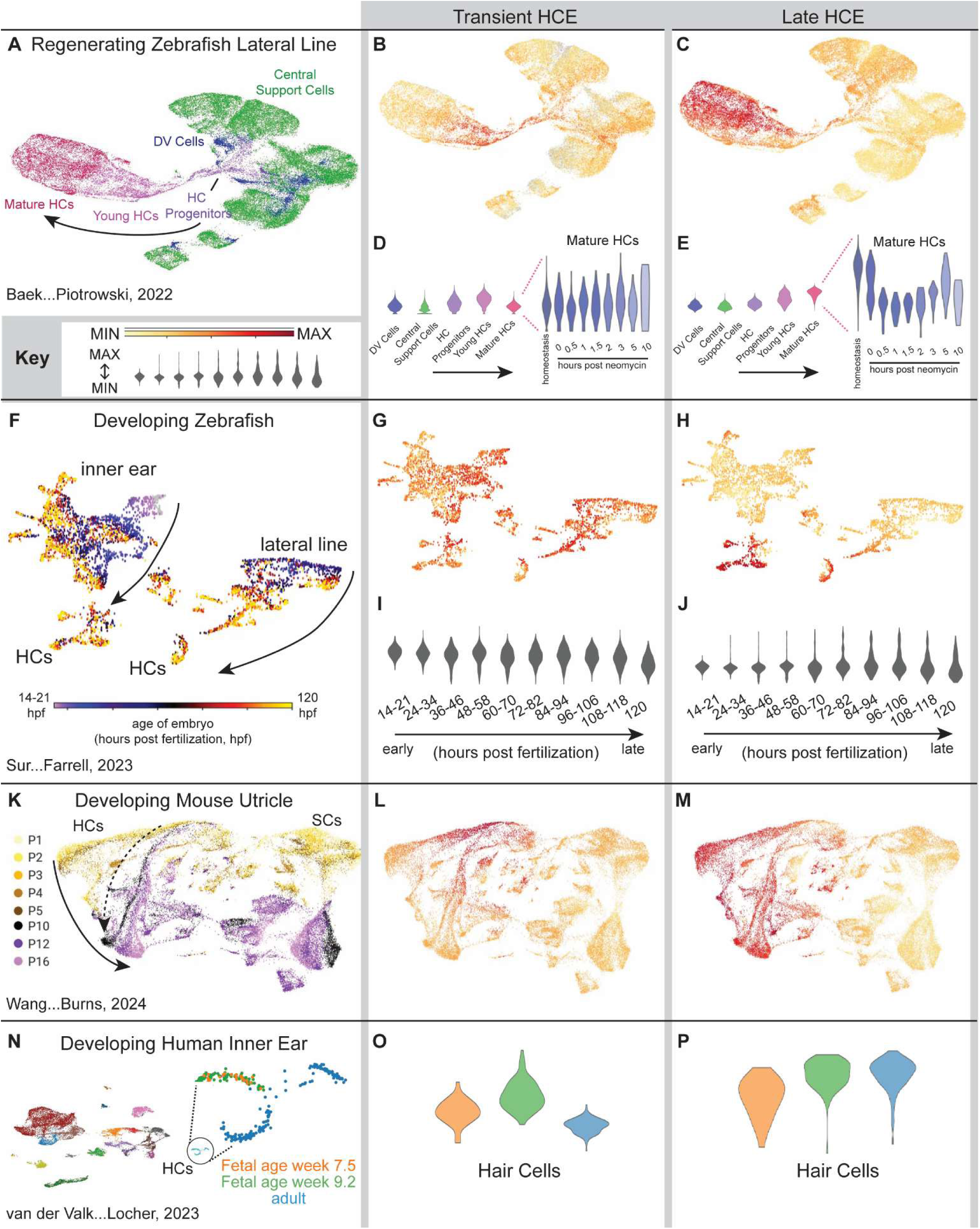
Transient and Late HCE genes show distinct transcriptomic dynamics in regeneration and development across species. (**A-E**) Transcriptomic analysis of the zebrafish lateral line in homeostasis and at intervals following neomycin-induced hair cell death revealed support cells (central and DV) and hair cells at various stages of maturity (progenitors, young, mature) (A). The Transient HCE (T-HCE) gene set shows strong expression in HC progenitors and young HCs (B) whereas the Late HCE (L-HCE) gene set is strongest in mature HCs (C), shown also by violin plots (left panels, D,E). When examined in mature HCs across the time course of regeneration (right panels, D,E), T-HCE gene expression changes little, but the L-HCE genes show a strong downregulation that starts to rebound 2 hours after neomycin treatment. (**F-J**) Inner ear and lateral line cells were subsetted from an atlas of the developing zebrafish and color coded based on the age of the larvae, from early (i.e. 14-21 hours post fertilization, hpf) to late (120 hpf). Two parallel trajectories can be inferred, resulting in discrete HC clusters (HCs). T-HCE genes (G,I) are broadly expressed, with slight downregulation in cells from the oldest larvae, shown as a feature plot (G) and in violin plots for each age (I). By contrast, L-HCE genes are enriched in both inner ear and lateral line HCs (H) and the signal increases as development proceeds (J). (**K-M**) UMAP analysis of single cells in developing mouse utricles (color coded by age, K) segregates HCs and support cells (SCs). Accordingly, T-HCE (L) and L-HCE (M) gene sets are both enriched in the HC clusters compared to SC clusters, with T-HCE genes more highly expressed at the beginning of the apparent trajectory and L-HCE genes progressively enriched along and at the end of the trajectory. Because HC developmental processes persist in the postnatal utricle due to homeostatic HC turnover, tissue from later stages contains the cellular trajectory of HC maturity in parallel to the trajectory of developing HCs in the early stages. This trajectory is indicated by the dashed arrow, while the developmental trajectory spanning animal ages is indicated by the solid arrow. (**N-P**) RNA from cells of the human fetal inner ear was sequenced a week 7.5 and week 9.2 and compared to cells of the adult inner ear. Analysis of the HCs from each age revealed a peak of T-HCE expression at week 9.2 and much lower expression in adults. Conversely, the L-HCE signal increased from week 7.5 to 9.2 and remained high in adult HCs. Raw data from relevant cell types were taken from the indicated publications and reanalyzed on gEAR, so the UMAPs are not identical to what was previously published. Key applies to heat maps in B, C, G, H, L, and M and to violin plots in D, E, I, J, O, and P.

Collectively, these analyses illustrate that the pan-HCE gene list offers a reliable indicator of development, regeneration, and reprogramming that can be shared across the field to query hair cell identity and maturity in data across species.

### A tool to query hair cell gene expression across species

Although we focused on identifying genes shared by most hair cells, the aggregate dataset can be used to query hair cells in multiple ways. For instance, researchers seeking markers for young mouse hair cells might want to identify genes with high AUCs and also high expression levels in mouse hair cells, regardless of how those genes are expressed in other species. Additionally, all AUC values over 0.5 indicate enrichment but lower scores indicate less enrichment (i.e. *OTOF* vs *GPX2*, **Fig. 2**) and there may be times when different AUC thresholds are appropriate, for example set at a high stringency by the user (**Fig. 3**) or slightly lower stringency using machine learning (**Fig. 4**). At other times, a zebrafish or chicken researcher may be interested in how mammalian hair cells express a gene they have found to be important for hair cell regeneration. To facilitate additional custom comparisons, we used R Shiny to create the HRP Hair Cell Gene Explorer, a publicly available, web-based data exploration environment (https://herblab.shinyapps.io/shinyapp/) that is integrated with a profile on the gEAR portal (Orvis et al., 2021), which collectively provide extensive capabilities to explore the expression patterns of hair cell-enriched genes and perform custom analyses (**Fig. 8**).

**Figure 8:**
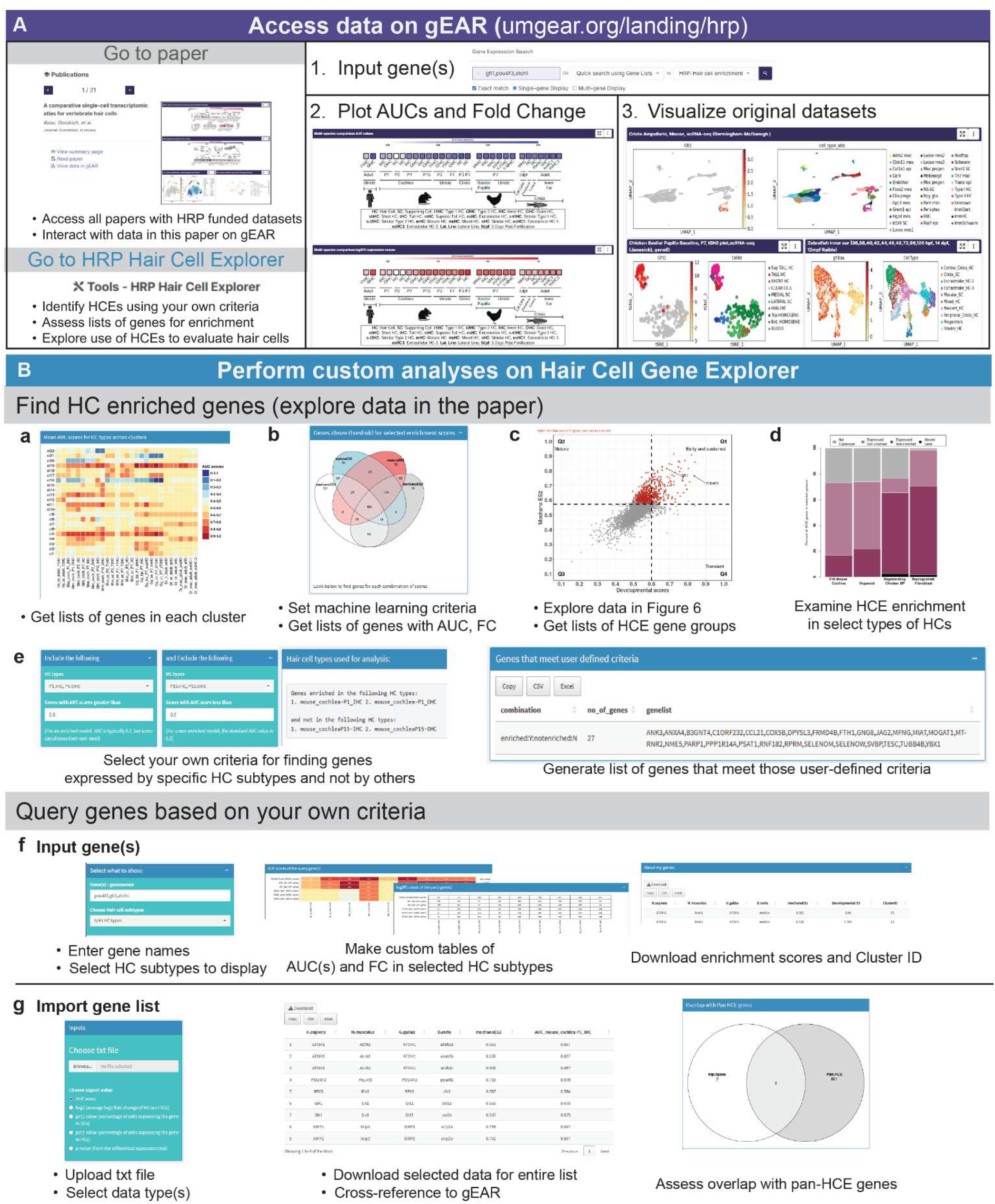
A public resource for evaluating hair cell gene expression. (**A**) Data in this manuscript can be accessed freely through the Hearing Restoration Project (HRP) landing page on the gEAR portal (https://umgear.org/landing/hrp/index.html). Through this page, users can input genes of interest (1.), recover AUCs and Fold Change (2.), and visualize the original gene expression data in each dataset (3.). Alternatively, users can route to the HRP Hair Cell Gene Explorer, which enables custom analyses (https://herblab.shinyapps.io/shinyapp/). (**B**) The HRP Hair Cell Gene Explorer is a Shiny App tool that makes it easy to create lists of HCE genes that meet certain criteria, such as cluster ID (a) or custom machine learning parameters (b). Users can also interact with the data in Figure 6 by changing thresholds and/or scoring method(s) (c) and by plotting the proportions of genes in each HCE group that are expressed and/or enriched in the example datasets (d). Users can identify HCE genes that meet specific criteria, such as expression in young but not mature mouse hair cells (e). Genes of interest can be inputted to produce custom heatmaps and tables of information about their expression and enrichment (f). Alternatively, users can import a gene list and gather information about AUC, log_2_ fold change, the percentage of cells in each hair cell (pct1) or supporting cell (pct2) cluster that express the gene and the p value. This function also identifies which genes are pan-HCE genes defined in this paper (g). Expression of all or one of the subsets of HCE genes can be projected onto any dataset of interest using the ProjectR tool in gEAR, as in Fig. 7.

The HRP Hair Cell Gene Explorer allows users to screen differential expression statistics across all conditions for genes that meet their criteria for AUC values, fold change between hair cells and supporting cells, and the percentage of supporting cells that have any detectable expression, as well as which species and hair cell types are included. Additionally, gene lists that are generated by each query are automatically linked to the original datasets on gEAR so that users can directly evaluate expression across all cell types in the dataset. Users can also explore the expression patterns of gene sets defined by our analyses, including the gene co-expression clusters (**Fig. 3**) and HCE genes (**Fig. 4**) reported here. Further, this resource offers an accessible way to quickly examine how any gene of interest behaves in all 43 hair cell populations in our analysis, including the degree of enrichment relative to supporting cells. This is a useful way for any researcher to gauge whether a gene they identified as key for hair cell development, regeneration, or function in their system is expressed as predicted in other organs or species. An additional feature allows users to upload lists of genes, for instance from their own RNA sequencing experiments, to identify those that are normally enriched in hair cells relative to supporting cells, as well as the numbers of T-HCE, ES-HCE, and L-HCE genes that are expressed. Each of these gene sets can also be projected onto a dataset of interest to identify hair cells, reveal relevant dynamics in the population, and directly compare the effects of experimental manipulations in a quantitative manner, as we did to assess T- and L- HCE genes in scRNA-sequencing datasets from zebrafish, mouse, and human (**Fig. 7**). The HRP Hair Cell Gene Explorer lowers the barrier for making comparisons across developmental stages, organs, and species, facilitating future investigation of similar *vs.* unique mechanisms of hair cell differentiation and function.

## Discussion

Although gene expression programs for hair cells have been described for many developmental stages, tissues, and species, it has been difficult to identify shared or unique modules and motifs as few cross-study comparisons have been performed (Wu et al., 2023)(Giffen et al., 2024) (Shi et al., 2024). Here, we developed an approach to compare data even when they come from different species or were generated using different methods. We first linked each gene to all predicted orthologs in other species, enabling comparisons beyond one-to-one orthologs. Then, rather than directly integrating datasets, which can obscure biological differences by forcing cell types to cluster on the basis of a small number of genes, we identified and meta-analyzed sets of genes enriched in each hair cell cluster relative to supporting cells in the same organ. By focusing not on expression levels but instead on the probability that a gene distinguishes two cell types, this approach enabled us to make direct comparisons that were robust to methodological differences, while also informing the biologically relevant relationship between supporting cells and hair cells. Unbiased hierarchical clustering and machine learning approaches defined a large set of hair cell-enriched (HCE) genes, including almost 1000 genes that are expressed at higher levels in hair cells than supporting cells across all organs and species examined. Known hair cell markers were on the list, and independent *in situ* hybridization and spatial transcriptomics for a subset of predicted pan-HCE genes further validated the approach. We also showed that this panel of pan-HCE genes can be used to assess the identity and degree of maturity of hair cells in independent transcriptomic datasets from different species, without calculating enrichment scores. Thus, this cross-species HCE database offers a useful and reliable framework for comparing and contrasting the genes that make hair cells different from supporting cells.

Examining hair cells across multiple organs and species identified hundreds of genes that are enriched in nearly all hair cells relative to supporting cells, strongly supporting the hypothesis that hair cells with dramatically different morphologies and functions rely on a shared molecular foundation. Many pan-HCE genes encode known stereociliary proteins, consistent with the presence of stereociliary bundles on the apical surface of all hair cells but not supporting cells. There was also a notable association with synaptic biology: This included known players such as VGLUT3 and Cav1.3, but also APBA1/MINT1, SYT14, and SNAP91, which are associated with synaptic vesicles and could contribute to ribbon synapse properties (Obholzer et al., 2008)(Sheets et al., 2012)(Kroll et al., 2020)(Okamoto and Südhof, 1997)(Ho et al., 2003)(Fukuda and Mikoshiba, 2001). These findings confirm and extend previous cross-species comparisons, which revealed shared use of genes associated with stereociliary bundles and synaptic transmission (Giffen et al., 2024) but relied on fewer species and developmental ages or did not involve direct single cell comparisons (Wang et al., 2024b)(Shi et al., 2024)(Wu et al., 2023)(Giffen et al., 2024). Future investigation of pan-HCE genes will clarify common stereociliary and synaptic mechanisms across hair cell types.

In addition to this shared core module, the precise sets of HCE genes varied widely among species, as demonstrated by unsupervised hierarchical clustering (**Fig. 3**). This may reflect flexibility across taxa in which genes are used to create features such as the stereociliary bundle or ribbon synapse. This variation may also indicate that different gene expression programs underlie unique aspects of hair cell biology in specific subtypes, organs, or species. Notably, MetaNeighbor analysis suggested that hair cells within a species rely on similar gene cohorts, with only weak correlations across hair cells from different species, even when they are from the same organ. We also did not find any clear homolog for the OHC in non-mammalian species. The apparent lack of homology among mature hair cell types stands in contrast to obvious cellular homologies in the retina from different species, where molecular similarity was enough to identify a type of retinal ganglion cell in mice that was previously thought to be present only in primates (Hahn et al., 2023). Further investigation of the 22 distinct co-regulated gene clusters identified here is needed to fully understand the biological implications of the variation among hair cells.

The ∼1000 genes reliably enriched across most or all hair cell subtypes offer a more powerful basis for comparison than canonical markers such as Myo7a. One advantage is that they capture expression dynamics relative to supporting cells. They also provide a quantitative measure of maturity and how “hair cell-like” a reprogrammed or regenerated cell is. By grounding the analysis in hair cell/supporting cell differences, which emerge both during development and in regeneration, our approach has the potential to reveal insight into the gene regulatory events that drive hair cell specification, diversification, and maturation. Indeed, the broader collection of HCE genes contains many transcription factors, including canonical regulators of hair cell identity such as Atoh1, Gfi1, and Pou4f3, but also less characterized transcription factors such as Lhx3 (Hertzano et al., 2007). Detailed analysis of additional transcription factors, whether shared or subtype-specific, will help establish a comprehensive approach for directing cells toward a mature hair cell phenotype, which is a prerequisite for functional regeneration.

Although we have focused on shared HCE genes, our data can also be used to ask a wide range of questions about hair cells. To facilitate customized exploration, all data and analyses are accessible through an online portal (gEAR), which hosts the HRP Hair Cell Gene Explorer tool. Users can query genes enriched across any subset of the hair cell groups examined, such as all mammalian hair cells, all mouse hair cells, or vestibular hair cells specifically. These kinds of analyses could potentially facilitate the design of cell-type specific tools, aid in the interpretation of genetic variation associated with deafness (Kalra et al., 2020), and advance basic understanding of hair cell function. Researchers can project the entire pan-HCE gene set or just the T-HCE and L-HCE gene sets onto their own data to evaluate how newly produced hair cells mature following specific manipulations, potentially revealing genes resistant to reprogramming and identifying developmental bottlenecks. This approach also enables comparison to other datasets hosted on the gEAR portal. The HCE gene list was generated using stringent criteria, so important players may not be included; however, the enrichment of any gene relative to supporting cells can still be examined through this resource, and users can adjust AUC thresholds to suit their analyses.

Several limitations should be considered. AUC-based enrichment captures whether expression differs between cell types but not the magnitude of that difference; a gene enriched in hair cells may still be expressed in supporting cells, just at lower levels. Indeed, many pan-HCE genes are expressed at low levels in zebrafish supporting cells, and whether this is biologically meaningful warrants future investigation. Conversely, genes expressed transiently or at low levels may have been missed due to low AUC scores, and genes with high AUC scores may also be expressed in other inner ear cell types such as neurons. These relationships can be explored through the HRP Hair Cell Gene Explorer and gEAR. In addition, a gene’s AUC score does not necessarily reflect its functional importance in hair cells, just its relative enrichment compared to supporting cells. Our analysis was constrained by scRNA-seq studies available when we started, so we were not able to include data from the human cochlea, and not all mouse vestibular organs are represented. Additionally, most of the 43 hair cell populations are from young animals and may not be fully mature. Subtle differences in the degree of maturity may contribute to some of the clustering results, for instance. More temporal data are needed to clarify how early hair cell development is initiated across species. Our analyses also reflect baseline conditions; key differences among species may only become apparent upon perturbation. Similarly, some of the mouse/human differences may be due to responses of the human utricle to surgical extraction and tissue dissociation. Finally, supporting cells were pooled in this analysis, and subtype-level differences may have been obscured. A similar analysis of supporting cells across species, organs, and time points would provide a valuable extension of this resource.

We hope that the findings reported here will prove useful to inform work across the field. The HCE gene set provides a strong framework for deducing the trajectories by which supporting cells give rise to well-differentiated hair cells in different species, to unveil gene regulatory mechanisms that differ between regenerative and non-regenerative conditions, and to benchmark outcomes for gene therapeutic strategies designed to restore hearing.

## Materials and Methods

### Experimental Design

We analyzed a comprehensive collection of previously generated scRNA-seq datasets profiling mechanosensory hair cells and supporting cells under homeostatic conditions. We selected available datasets spanning diverse hearing and vestibular organs from developmental through young adult timepoints. We included datasets from vertebrate clades that differ in the capacity of hair cells to regenerate after injury, including humans, mice, chickens, and zebrafish. We excluded non-homeostatic conditions, such as datasets profiling the responses to chemical or noise-induced damage to hair cells, as well as genetic knockout experiments. Most of the scRNA-seq datasets analyzed were produced using the 10x Genomics platform, which uses droplet microfluidics to generate high-throughput transcriptomic data from thousands of single cells in parallel. A smaller subset of datasets was produced using SMART-seq, a full-length transcript scRNA-seq method that provides more comprehensive coverage of transcript isoforms. Our analysis started from processed cell x gene counts matrices and associated cell type annotations from scRNA-seq experiments, obtained from the authors of each study or public repositories. See **Table S1**.

### MetaNeighbor analysis

The MetaNeighbor (v1.22.0) function MetaNeighborUS (fast_version = TRUE) run in R (v4.3.0) was used to compare hair cell populations across multiple species (Crow et al., 2018). Each dataset was subsetted to remove all populations other than hair cells and downsampled to no more than 100 cells of each subtype. The top 2000 most variable genes were identified within the resulting dataset using Seurat, and those genes were used in the MetaNeighbor calculation. Correlation scores calculated for all pairs of hair cell populations were represented in a heat map.

### Annotation of orthologous genes across species

Genes from mouse, chicken, and zebrafish were mapped to their human orthologs, based on the union of annotations in the Alliance, JAX, and Ensembl (BioMart) databases. If a human gene was mapped to 2-4 orthologs in another species, we included all of the mapped orthologs. However, human genes mapped to >4 orthologs in other species were removed from the analysis. See **Table S2** for correspondences.

### Calculation of HCE genes in each condition

We assessed gene enrichment in individual hair cell subtypes relative to supporting cells. To ensure consistency across datasets, we used the area under the receiver operating characteristic curve (AUC) to quantify gene enrichment, using Seurat FindMarkers() with test.use = ‘roc’ (Butler et al., 2018). This metric evaluates a gene’s ability to distinguish hair cells from supporting cells within each dataset. An AUC score of 1 indicates perfect discrimination, where the gene is exclusively expressed in hair cells or supporting cells, whereas an AUC score of 0.5 suggests no discriminatory power between the two cell types. To distinguish between hair cells and supporting cells, we first set AUC scores below 0.5 to 0.5. Then, for the remaining AUC scores, genes enriched in supporting cells relative to hair cells were defined as those with log_2_ fold change [logFC] < 0; for these genes, we reassigned the AUC score as 1 – AUC. This transformation ensures that genes with AUC > 0.5 are interpreted as enriched in hair cells, while those with AUC < 0.5 are enriched in supporting cells. In datasets that included multiple supporting cell types, we grouped all SC types into a single category labeled “supporting cell”, which served as the reference group for enrichment analysis across each hair cell subtype. We constructed a combined matrix in which rows represent genes and columns represent hair cell subtypes. Each cell in this matrix contains the computed AUC score, log_2_FC, and p-value, summarizing the degree of enrichment for each gene in the corresponding hair cell subtype. See **Table S2**.

### Clustering of genes based on their enrichment patterns across conditions

To analyze gene expression patterns across hair cell subtypes, we focused on the 29 mechanotransduction-capable hair cell subtypes. Genes enriched in hair cells were selected based on an AUC score of ≥0.7 in at least one cell type, resulting in 4,964 genes used for clustering. Unsupervised hierarchical clustering was performed using the Ward.D2 method on a Euclidean distance matrix derived from the genes’ AUC scores across the 29 cell types (**Figure 3–figure supplement 1**). To refine the clusters, genes with low correlation to their cluster centroids were removed, and highly similar clusters were merged. This process yielded 22 non-overlapping gene clusters. Module membership for each gene was then defined by its correlation with the cluster centroid’s AUC profile. In addition, to assess the overall AUC score patterns across all hair cell subtypes, we calculated the average AUC score of all genes within each cluster, specific to each subtype (**Fig. 3**).

### Machine-learning approach to derive pan-HCE genes

Four alternative approaches were considered to produce aggregate scores for each gene across hair cell subtypes, including directly summing the AUC scores (enrichment score [ES]1) vs. weighting the scores to account for uneven representation of hair cell types across species (14 mouse, 6 chicken, 7 zebrafish, and 2 human; ES2), as well as summing the AUC scores for all 29 hair cell populations capable of mechanotransduction vs. the 23 most mature populations (as in **Fig. 1C**). These four approaches produce four scores: ES1 and ES2 for all and ‘mechano’ subtypes for each of the genes, providing a rank-ordering of all genes from the most consistently and highly enriched in hair cells to the most highly enriched in supporting cells. In principle, this rank-ordered list could be cut at any level depending on the desired use. For instance, some analyses require just a small number of very strongly hair cell-enriched (HCE) genes, and we recommend the top 50 genes for this purpose. However, this set excludes many genes known to be enriched in hair cells. To define a more comprehensive set of HCE genes, we compared our rank-ordered gene list to “gold standard” genes that have previously been shown to be enriched in hair cells by independent approaches. Specifically, we defined our gold standards as the union of hair cell enriched genes from five sources: (i) translatome profiling of hair cells in the mouse cochlea at P6 (Tao et al., 2021) or P28 (Liu et al., 2018); (ii) RNA-seq of FACS-sorted hair cells in the chicken utricle at P7 (Scheibinger et al., 2022); (iii) mass spectrometry proteomics of hair bundles in the chicken and mouse inner ear (Krey and Barr-Gillespie, 2019)(Wilmarth et al., 2015); (iv) ontology-based annotations of hair cell genes from the Zebrafish Information Network (ZFIN)(Sprague et al., 2003); (v) a manually curated list of hair cell-enriched genes produced by co-authors of this study who were blinded to our scRNA-seq-defined gene lists (**Table S4**). These true-positive HCE genes were at the top of our ranked list, but some true positives had lower ranks. At each rank threshold, we calculated the precision -- true positives / [true positives + false positives] – and recall – true positives / [true positives + false negatives]. At more lenient thresholds, the precision decreases (more false positives), but the recall improves (more true positives). We note that a “false positive” in this case merely indicates that a gene was not detected as HCE in the independent studies used to define the gold standards – many of these novel findings may in fact be real HCE genes. As an optimal threshold, we used the precision-recall breakeven point, which is the threshold at which precision is equal to recall and is considered a good threshold when one desires to balance the rates of true positives and false negatives. The precision and recall curves were calculated separately for each of the four ranking strategies. The four resulting lists were very similar, indicating that our approach is reasonably robust to these parameters. The final list of HCE genes was defined as the intersection among the four gene lists.

### Gene set enrichment analysis

Gene Set Enrichment Analysis (GSEA) was conducted on hair cell-enriched genes using their AUC scores. The analysis was performed with the “gseGO” function from the clusterProfiler R package (Wu et al., 2021). Among all significantly enriched Biological Process (BP), Molecular Function (MF), and Cellular Component (CC) Gene Ontology (GO) terms identified, independent terms were retained based on gene overlap. Enrichment analyses of custom gene lists were performed using Fisher’s exact tests. Known deafness-associated genes were obtained from the Hereditary Hearing Loss Homepage (Walls WD, Azaiez H, Smith RJH., n.d.). Stereocilia components were compiled from two studies that performed proteomics of stereocilia of chicken (Shin et al., 2013) and mouse hair cells (ProteomeXchange PXD002167) (*55*), respectively. GO enrichments for the genes in each co-expression cluster were calculated using the enrichGO function from the clusterProfiler package (**Table S3,S6**).

### In situ hybridization methods – RNAscope, HCR, and Spatial Transcriptomics

#### Mouse RNAScope

One-month-old C57BL/6 mice were euthanized. Temporal bones were extracted and incubated overnight in RNase-free 4% paraformaldehyde for fixation followed by 3 days in 150 mM EDTA at 4°C for decalcification. The samples were then incubated in 30% sucrose followed by incubation in a 1:1 mix of 30% sucrose and Super Cryoembedding Medium freezing medium (SECTION-LAB Co. Ltd.) for 3 hours and flash frozen in liquid nitrogen. The samples were kept at -80°C until processing. The tissue was sectioned at 12 μm using a CM 1850 cryostat (Leica). *In-situ* hybridization was performed following Advanced Cell Diagnostics’ (ACD) instructions for the RNAscope Multiplex Fluorescent Reagent Kit v2 with the following ACD pre-designed probes: Mm-Apba1, Mm-Syt14, Mm-Rab3ip, Mm- Runx1t1, Mm-Snap91, Mm-Pgm2l1, Mm-Pcsk5, Mm-Fscn2, Mm-Tmem255b and Mm-Xirp2. Images were acquired with a Nikon AX R Confocal Microscope System.

#### Wholemount mouse and zebrafish HCR

Hybridization chain reaction (HCR) *in situ* hybridizations (Molecular Instruments, HCR v3.0) were performed in whole-mounts of adult mouse utricles and whole larval zebrafish as described previously (Choi et al., 2016)(Beaulieu et al., 2024). After euthanasia, fish were fixed in 4% PFA at 4°C for 18-48 h, and mouse temporal bones were fixed in 4% PFA for 2-4 hours at room temperature. Tissues were dissected, washed with PBS and transferred to methanol for storage at −20°C. Samples (fish or mouse utricles) were rehydrated using a gradation of methanol and PBS containing 0.1% Tween20, treated with 10 μg/ml proteinase K for 25 min and post-fixed with 4% PFA for 20 min at room temperature. Probes were designed by Molecular Instruments. Samples were pre-hybridized with a probe hybridization buffer for 30 min at 37°C, then incubated with probes overnight at 37°C. Samples were washed with 5×SSCT to remove excess probes. For the amplification stage, samples were pre-incubated with an amplification buffer for 30 min at room temperature and incubated with hairpins overnight in the dark at room temperature. Excess hairpins were removed by washing with 5×SSCT, and samples were transferred to storage buffer and kept in the dark at 4°C until imaging. Mouse utricles were counterlabeled with antibodies to Calb2 (Santa Cruz) to label type II hair cells or calyx afferents. Confocal images were captured using a Zeiss LSM-980 with Airyscan 2.0 or an Olympus Fluoview 1000. Z-stacks for fish were taken using a 25×/0.8 water objective at intervals of 0.58 μm, and for mouse using 60x objective with intervals of 1 µm. Airyscan processing was performed at standard strength using Zen Blue software (Zeiss, www.zeiss.com). Olympus file processing was facilitated by Fiji (<u>imagej.net</u>)

#### Mouse Spatial Transcriptomics

Inner ears were harvested from CBA/J mice at 4 weeks-of-age and fixed overnight in 4% paraformaldehyde and 1X phosphate buffer solution. Tissue was dehydrated in ethanol, cleared using xylene, and embedded in paraffin with a Leica Histocore Pegasus paraffin processor. Embedded tissue was sectioned at a thickness of 5 μm and placed on Fisherbrand^TM^ Superfrost^TM^ Plus microscope slides. Tissue slides were stained with hematoxylin and eosin. Stained tissue was imaged at 40x using a Zeiss Axioscan 7, courtesy of the NICHD Microscopy and Imaging Core. The standard 10x Genomics Visium HD platform and protocol was applied to tissue slides for RNA capture and cDNA library preparation. Samples were sequenced by the Genomics and Computational Biology Core using an Illumina HiSeq1500. Image data was aligned with transcriptomics data by following the standard 10x Genomics Space Ranger pipeline. The 10x Genomics Loupe Browser software was used for cell annotation and analysis of gene expression levels.

#### Chicken HCR

*In situ* mRNA detection by hybridization chain reaction (HCR) was performed in chicken basilar papilla and utricle using RNase-free reagents and following established protocols(Sato et al., 2024b). Briefly, inner ear organs were isolated under sterile, RNase-free conditions, fixed in 4% paraformaldehyde in PBS, sectioned on a Leica vibratome into 70 µm slices, equilibrated in 50% methanol for 10 min, then transferred to 100% methanol and stored at −20°C. For hybridization, up to eight sections were transferred from methanol and mounted on Gold Seal™ UltraStick™ Adhesion Microscope Slides (Thermo Scientific) to ensure firm attachment during subsequent steps. Hybridization chain reaction (HCR v3.0, Molecular Instruments) was performed according to the manufacturer’s protocol with previously described modifications (Choi et al., 2018)(Sato et al., 2024b). Sections were rehydrated, permeabilized with proteinase K (10 µg/mL, 10 min, room temperature), equilibrated in hybridization buffer, and incubated overnight at 37 °C with probe sets targeting the genes of interest. The next day, excess probes were removed with a series of stringent, pre-warmed washes, followed by equilibration in amplification buffer. For signal amplification, fluorescent DNA hairpins (H1 and H2) were snap-cooled (95 °C for 90 s, then 30 min at room temperature in the dark) and applied in amplification buffer for 16– 18 h at room temperature. After washing, sections were counterstained with DAPI (1 µg/mL) and mounted. Negative controls (no-probe and scrambled-probe conditions) were processed in parallel to assess specificity. Imaging was performed on a Zeiss LSM 700 confocal microscope with identical acquisition settings across samples.

#### Statistical Analysis

**Statistical** analyses compared gene expression in each hair cell subtype to supporting cells from the same condition. Hair cell enriched genes were clustered and ranked based on the level of enrichment across hair cell subtypes. Equivalent analyses were performed to compare mechanosensation-capable hair cells to developing and regenerating hair cells. Detailed descriptions of the statistical approaches are presented in the preceding methods subsections.

## Acknowledgments

We thank Chris Gralapp (chrisgralapp.com) for her contributions to the scientific illustrations. We are grateful to Timothy Higdon, Christopher Geissler, and Yishane Lee of the Hearing Health Foundation for their tireless efforts to support the HRP consortium.

## Funding

Hearing Health Foundation’s Hearing Restoration Project

National Institutes of Health grant DC021755 (NB)

Canadian Institutes of Health Research – CIHR PJT-183579 (EL, AD)

Michael and Sonja Koerner Charitable Foundation (AD)

National Institutes of Health R01 DC020322 (AE)

National Institutes of Health R01 DC020268 (KG)

National Institutes of Health R01 DC014832 (AG)

National Institutes of Health R01 DC019619 (SH)

Intramural Research Program, National Institutes of Health ZIA DC000094 (RH)

National Institutes of Health U01 DC019370 (AM, RH)

National Institutes of Health R01 DC015488 (TP)

Stowers Institute for Medical Research (TP)

National Institutes of Health CoBRE RPL grant P20GM139762-RPL (LT)

National Institutes of Health R21 DC020773 (LT)

State of Nebraska LB692 (LT)

National Institutes of Health R01 DC006283 (MW)

This research was supported in part by the Intramural Research Program of the National Institutes of Health (NIH). The contributions of the NIH author(s) are considered Works of the United States Government. The findings and conclusions presented in this paper are those of the author(s) and do not necessarily reflect the views of the NIH or the U.S. Department of Health and Human Services.

## Notes

### Competing Interest Statement

The authors have declared no competing interest.

## References

Baek S, Tran NTT, Diaz DC, Tsai Y-Y, Navajas Acedo J, Lush ME, Piotrowski T. 2022. Single-cell transcriptome analysis reveals three sequential phases of gene expression during zebrafish sensory hair cell regeneration. Developmental Cell 57:799–819.e6. DOI: 10.1016/j.devcel.2022.03.001, PMID: 35316618

Beaulieu MO, Thomas ED, Raible DW. 2024. Transdifferentiation is temporally uncoupled from progenitor pool expansion during hair cell regeneration in the zebrafish inner ear. Development (Cambridge, England) 151:dev202944. DOI: 10.1242/dev.202944, PMID: 39045613

Benkafadar N, Sato MP, Ling AH, Janesick A, Scheibinger M, Jan TA, Heller S. 2024. An essential signaling cascade for avian auditory hair cell regeneration. Developmental Cell 59:280–291.e5. DOI: 10.1016/j.devcel.2023.11.028, PMID: 38128539

Bleckmann H, Zelick R. 2009. Lateral line system of fish. Integrative Zoology 4:13–25. DOI: 10.1111/j.1749-4877.2008.00131.x, PMID: 21392273

Bolz H, von Brederlow B, Ramírez A, Bryda EC, Kutsche K, Nothwang HG, Seeliger M, del C-Salcedó Cabrera M, Vila MC, Molina OP, Gal A, Kubisch C. 2001. Mutation of CDH23, encoding a new member of the cadherin gene family, causes Usher syndrome type 1D. Nature Genetics 27:108–112. DOI: 10.1038/83667, PMID: 11138009

Bork JM, Peters LM, Riazuddin S, Bernstein SL, Ahmed ZM, Ness SL, Polomeno R, Ramesh A, Schloss M, Srisailpathy CR, Wayne S, Bellman S, Desmukh D, Ahmed Z, Khan SN, Kaloustian VM, Li XC, Lalwani A, Riazuddin S, Bitner-Glindzicz M, Nance WE, Liu XZ, Wistow G, Smith RJ, Griffith AJ, Wilcox ER, Friedman TB, Morell RJ. 2001. Usher syndrome 1D and nonsyndromic autosomal recessive deafness DFNB12 are caused by allelic mutations of the novel cadherin-like gene CDH23. American Journal of Human Genetics 68:26–37. DOI: 10.1086/316954, PMID: 11090341

Brignull HR, Raible DW, Stone JS. 2009. Feathers and fins: non-mammalian models for hair cell regeneration. Brain Research 1277:12–23. DOI: 10.1016/j.brainres.2009.02.028, PMID: 19245801

Butler A, Hoffman P, Smibert P, Papalexi E, Satija R. 2018. Integrating single-cell transcriptomic data across different conditions, technologies, and species. Nature Biotechnology 36:411–420. DOI: 10.1038/nbt.4096, PMID: 29608179

Cai T, Groves AK. 2015. The Role of Atonal Factors in Mechanosensory Cell Specification and Function. Molecular Neurobiology 52:1315–1329. DOI: 10.1007/s12035-014-8925-0, PMID: 25339580

Chitramuthu BP, Baranowski DC, Cadieux B, Rousselet E, Seidah NG, Bennett HPJ. 2010. Molecular cloning and embryonic expression of zebrafish PCSK5 co-orthologues: functional assessment during lateral line development. Developmental Dynamics: An Official Publication of the American Association of Anatomists 239:2933–2946. DOI: 10.1002/dvdy.22426, PMID: 20882679

Choi HMT, Calvert CR, Husain N, Huss D, Barsi JC, Deverman BE, Hunter RC, Kato M, Lee SM, Abelin ACT, Rosenthal AZ, Akbari OS, Li Y, Hay BA, Sternberg PW, Patterson PH, Davidson EH, Mazmanian SK, Prober DA, van de Rijn M, Leadbetter JR, Newman DK, Readhead C, Bronner ME, Wold B, Lansford R, Sauka-Spengler T, Fraser SE, Pierce NA. 2016. Mapping a multiplexed zoo of mRNA expression. Development (Cambridge, England) 143:3632–3637. DOI: 10.1242/dev.140137, PMID: 27702788

Choi HMT, Schwarzkopf M, Fornace ME, Acharya A, Artavanis G, Stegmaier J, Cunha A, Pierce NA. 2018. Third-generation in situ hybridization chain reaction: multiplexed, quantitative, sensitive, versatile, robust. Development (Cambridge, England) 145:dev165753. DOI: 10.1242/dev.165753, PMID: 29945988

Chu FF, Doroshow JH, Esworthy RS. 1993. Expression, characterization, and tissue distribution of a new cellular selenium-dependent glutathione peroxidase, GSHPx-GI. The Journal of Biological Chemistry 268:2571– 2576. PMID: 8428933

Crow M, Paul A, Ballouz S, Huang ZJ, Gillis J. 2018. Characterizing the replicability of cell types defined by single cell RNA-sequencing data using MetaNeighbor. Nature Communications 9:884. DOI: 10.1038/s41467-018-03282-0, PMID: 29491377

Dallos P, Wu X, Cheatham MA, Gao J, Zheng J, Anderson CT, Jia S, Wang X, Cheng WHY, Sengupta S, He DZZ, Zuo J. 2008. Prestin-based outer hair cell motility is necessary for mammalian cochlear amplification. Neuron 58:333–339. DOI: 10.1016/j.neuron.2008.02.028, PMID: 18466744

Erkman L, McEvilly RJ, Luo L, Ryan AK, Hooshmand F, O’Connell SM, Keithley EM, Rapaport DH, Ryan AF, Rosenfeld MG. 1996. Role of transcription factors Brn-3.1 and Brn-3.2 in auditory and visual system development. Nature 381:603–606. DOI: 10.1038/381603a0, PMID: 8637595

Fettiplace R, Hackney CM. 2006. The sensory and motor roles of auditory hair cells. Nature Reviews. Neuroscience 7:19–29. DOI: 10.1038/nrn1828, PMID: 16371947

Fukuda M, Mikoshiba K. 2001. Characterization of KIAA1427 protein as an atypical synaptotagmin (Syt XIII). The Biochemical Journal 354:249–257. DOI: 10.1042/0264-6021:3540249, PMID: 11171101

Géléoc GSG, Holt JR. 2003. Developmental acquisition of sensory transduction in hair cells of the mouse inner ear. Nature Neuroscience 6:1019–1020. DOI: 10.1038/nn1120, PMID: 12973354

Giffen KP, Liu H, Yamane KL, Li Y, Chen L, Kramer KL, Zallocchi M, He DZ. 2024. Molecular specializations underlying phenotypic differences in inner ear hair cells of zebrafish and mice. Frontiers in Neurology 15:1437558. DOI: 10.3389/fneur.2024.1437558, PMID: 39484049

Golub JS, Tong L, Ngyuen TB, Hume CR, Palmiter RD, Rubel EW, Stone JS. 2012. Hair cell replacement in adult mouse utricles after targeted ablation of hair cells with diphtheria toxin. The Journal of Neuroscience: The Official Journal of the Society for Neuroscience 32:15093–15105. DOI: 10.1523/JNEUROSCI.1709-12.2012, PMID: 23100430

Guo J-Y, Xu J-Y, Gong S-S, Wang G-P. 2024. Roles of supporting cells in the maintenance and regeneration of the damaged inner ear: A literature review. Journal of Otology 19:234–240. DOI: 10.1016/j.joto.2024.07.007, PMID: 39776546

Hahn J, Monavarfeshani A, Qiao M, Kao AH, Kölsch Y, Kumar A, Kunze VP, Rasys AM, Richardson R, Wekselblatt JB, Baier H, Lucas RJ, Li W, Meister M, Trachtenberg JT, Yan W, Peng Y-R, Sanes JR, Shekhar K. 2023. Evolution of neuronal cell classes and types in the vertebrate retina. Nature 624:415–424. DOI: 10.1038/s41586-023-06638-9, PMID: 38092908

Hertzano R, Dror AA, Montcouquiol M, Ahmed ZM, Ellsworth B, Camper S, Friedman TB, Kelley MW, Avraham KB. 2007. Lhx3, a LIM domain transcription factor, is regulated by Pou4f3 in the auditory but not in the vestibular system. The European Journal of Neuroscience 25:999–1005. DOI: 10.1111/j.1460-9568.2007.05332.x, PMID: 17331196

Ho A, Morishita W, Hammer RE, Malenka RC, Sudhof TC. 2003. A role for Mints in transmitter release: Mint 1 knockout mice exhibit impaired GABAergic synaptic transmission. Proceedings of the National Academy of Sciences of the United States of America 100:1409–1414. DOI: 10.1073/pnas.252774899, PMID: 12547917

Hudspeth AJ. 2008. Making an effort to listen: mechanical amplification in the ear. Neuron 59:530–545. DOI: 10.1016/j.neuron.2008.07.012, PMID: 18760690

Janesick A, Scheibinger M, Benkafadar N, Kirti S, Ellwanger DC, Heller S. 2021. Cell-type identity of the avian cochlea. Cell Reports 34:108900. DOI: 10.1016/j.celrep.2021.108900, PMID: 33761346

Janesick AS, Scheibinger M, Benkafadar N, Kirti S, Heller S. 2022. Avian auditory hair cell regeneration is accompanied by JAK/STAT-dependent expression of immune-related genes in supporting cells. Development (Cambridge, England) 149:dev200113. DOI: 10.1242/dev.200113, PMID: 35420675

Jones JM, Montcouquiol M, Dabdoub A, Woods C, Kelley MW. 2006. Inhibitors of differentiation and DNA binding (Ids) regulate Math1 and hair cell formation during the development of the organ of Corti. The Journal of Neuroscience: The Official Journal of the Society for Neuroscience 26:550–558. DOI: 10.1523/JNEUROSCI.3859-05.2006, PMID: 16407553

Kalra G, Lenz D, Abdul-Aziz D, Hanna C, Basu M, Herb BR, Colantuoni C, Milon B, Saxena M, Shetty AC, Hertzano R, Shivdasani RA, Ament SA, Edge ASB. 2023. Cochlear organoids reveal transcriptional programs of postnatal hair cell differentiation from supporting cells. Cell Reports 42:113421. DOI: 10.1016/j.celrep.2023.113421, PMID: 37952154

Kalra G, Milon B, Casella AM, Herb BR, Humphries E, Song Y, Rose KP, Hertzano R, Ament SA. 2020. Biological insights from multi-omic analysis of 31 genomic risk loci for adult hearing difficulty. PLoS genetics 16:e1009025. DOI: 10.1371/journal.pgen.1009025, PMID: 32986727

Kawamoto K, Izumikawa M, Beyer LA, Atkin GM, Raphael Y. 2009. Spontaneous hair cell regeneration in the mouse utricle following gentamicin ototoxicity. Hearing Research 247:17–26. DOI: 10.1016/j.heares.2008.08.010, PMID: 18809482

Khan S, Chang R. 2013. Anatomy of the vestibular system: a review. NeuroRehabilitation 32:437–443. DOI: 10.3233/NRE-130866, PMID: 23648598

Kolla L, Kelly MC, Mann ZF, Anaya-Rocha A, Ellis K, Lemons A, Palermo AT, So KS, Mays JC, Orvis J, Burns JC, Hertzano R, Driver EC, Kelley MW. 2020. Characterization of the development of the mouse cochlear epithelium at the single cell level. Nature Communications 11:2389. DOI: 10.1038/s41467-020-16113-y, PMID: 32404924

Krey JF, Barr-Gillespie PG. 2019. Molecular Composition of Vestibular Hair Bundles. Cold Spring Harbor Perspectives in Medicine 9:a033209. DOI: 10.1101/cshperspect.a033209, PMID: 29844221

Krey JF, Sherman NE, Jeffery ED, Choi D, Barr-Gillespie PG. 2015. The proteome of mouse vestibular hair bundles over development. Scientific Data 2:150047. DOI: 10.1038/sdata.2015.47, PMID: 26401315

Kroll J, Özçete ÖD, Jung S, Maritzen T, Milosevic I, Wichmann C, Moser T. 2020. AP180 promotes release site clearance and clathrin-dependent vesicle reformation in mouse cochlear inner hair cells. Journal of Cell Science 133:jcs236737. DOI: 10.1242/jcs.236737, PMID: 31843760

Lelli A, Asai Y, Forge A, Holt JR, Géléoc GSG. 2009. Tonotopic gradient in the developmental acquisition of sensory transduction in outer hair cells of the mouse cochlea. Journal of Neurophysiology 101:2961– 2973. DOI: 10.1152/jn.00136.2009, PMID: 19339464

Liberman MC, Gao J, He DZZ, Wu X, Jia S, Zuo J. 2002. Prestin is required for electromotility of the outer hair cell and for the cochlear amplifier. Nature 419:300–304. DOI: 10.1038/nature01059, PMID: 12239568

Liu H, Chen L, Giffen KP, Stringham ST, Li Y, Judge PD, Beisel KW, He DZZ. 2018. Cell-Specific Transcriptome Analysis Shows That Adult Pillar and Deiters’ Cells Express Genes Encoding Machinery for Specializations of Cochlear Hair Cells. Frontiers in Molecular Neuroscience 11:356. DOI: 10.3389/fnmol.2018.00356, PMID: 30327589

Luca E, Ibeh N, Yamamoto R, Liang W, Bennett D, Lin V, Chen J, Lovett M, Dabdoub A. 2025. Revealing heterogeneity and damage response in the adult human utricle. Nature Communications 17:9. DOI: 10.1038/s41467-025-66358-8, PMID: 41372202

McLean WJ, Yin X, Lu L, Lenz DR, McLean D, Langer R, Karp JM, Edge ASB. 2017. Clonal Expansion of Lgr5-Positive Cells from Mammalian Cochlea and High-Purity Generation of Sensory Hair Cells. Cell Reports 18:1917–1929. DOI: 10.1016/j.celrep.2017.01.066, PMID: 28228258

Menendez L, Trecek T, Gopalakrishnan S, Tao L, Markowitz AL, Yu HV, Wang X, Llamas J, Huang C, Lee J, Kalluri R, Ichida J, Segil N. 2020. Generation of inner ear hair cells by direct lineage conversion of primary somatic cells. eLife 9:e55249. DOI: 10.7554/eLife.55249, PMID: 32602462

Ó Maoiléidigh D, Ricci AJ. 2019. A Bundle of Mechanisms: Inner-Ear Hair-Cell Mechanotransduction. Trends in Neurosciences 42:221–236. DOI: 10.1016/j.tins.2018.12.006, PMID: 30661717

Obholzer N, Wolfson S, Trapani JG, Mo W, Nechiporuk A, Busch-Nentwich E, Seiler C, Sidi S, Söllner C, Duncan RN, Boehland A, Nicolson T. 2008. Vesicular glutamate transporter 3 is required for synaptic transmission in zebrafish hair cells. The Journal of Neuroscience: The Official Journal of the Society for Neuroscience 28:2110–2118. DOI: 10.1523/JNEUROSCI.5230-07.2008, PMID: 18305245

Okamoto M, Südhof TC. 1997. Mints, Munc18-interacting proteins in synaptic vesicle exocytosis. The Journal of Biological Chemistry 272:31459–31464. DOI: 10.1074/jbc.272.50.31459, PMID: 9395480

Orvis J, Gottfried B, Kancherla J, Adkins RS, Song Y, Dror AA, Olley D, Rose K, Chrysostomou E, Kelly MC, Milon B, Matern MS, Azaiez H, Herb B, Colantuoni C, Carter RL, Ament SA, Kelley MW, White O, Bravo HC, Mahurkar A, Hertzano R. 2021. gEAR: Gene Expression Analysis Resource portal for community-driven, multi-omic data exploration. Nature Methods 18:843–844. DOI: 10.1038/s41592-021-01200-9, PMID: 34172972

Pan B, Akyuz N, Liu X-P, Asai Y, Nist-Lund C, Kurima K, Derfler BH, György B, Limapichat W, Walujkar S, Wimalasena LN, Sotomayor M, Corey DP, Holt JR. 2018. TMC1 Forms the Pore of Mechanosensory Transduction Channels in Vertebrate Inner Ear Hair Cells. Neuron 99:736–753.e6. DOI: 10.1016/j.neuron.2018.07.033, PMID: 30138589

Pangršič T, Reisinger E, Moser T. 2012. Otoferlin: a multi-C2 domain protein essential for hearing. Trends in Neurosciences 35:671–680. DOI: 10.1016/j.tins.2012.08.002, PMID: 22959777

Popper AN, Fay RR. 1993. Sound detection and processing by fish: critical review and major research questions. Brain, Behavior and Evolution 41:14–38. DOI: 10.1159/000113821, PMID: 8431753

Pullin JM, McCarthy DJ. 2024. A comparison of marker gene selection methods for single-cell RNA sequencing data. Genome Biology 25:56. DOI: 10.1186/s13059-024-03183-0, PMID: 38409056

Qiu X, Müller U. 2022. Sensing sound: Cellular specializations and molecular force sensors. Neuron 110:3667– 3687. DOI: 10.1016/j.neuron.2022.09.018, PMID: 36223766

Ranum PT, Goodwin AT, Yoshimura H, Kolbe DL, Walls WD, Koh J-Y, He DZZ, Smith RJH. 2019. Insights into the Biology of Hearing and Deafness Revealed by Single-Cell RNA Sequencing. Cell Reports 26:3160–3171.e3. DOI: 10.1016/j.celrep.2019.02.053, PMID: 30865901

Sandler JE, Tsai Y-Y, Chen S, Sabin L, Lush ME, Sur A, Ellis E, Tran NTT, Cook M, Scott AR, Kniss JS, Farrell JA, Piotrowski T. 2025. prdm1a drives a fate switch between hair cells of different mechanosensory organs. Nature Communications 16:7662. DOI: 10.1038/s41467-025-62942-0, PMID: 40825768

Sato MP, Benkafadar N, Heller S. 2024a. Hair cell regeneration, reinnervation, and restoration of hearing thresholds in the avian hearing organ. Cell Reports 43:113822. DOI: 10.1016/j.celrep.2024.113822, PMID: 38393948

Sato MP, Huang AP, Heller S, Benkafadar N. 2024b. Protocol for in vivo elimination of avian auditory hair cells, multiplexed mRNA detection, immunohistochemistry, and S-phase labeling. STAR protocols 5:103118. DOI: 10.1016/j.xpro.2024.103118, PMID: 38852155

Scheffer DI, Zhang D-S, Shen J, Indzhykulian A, Karavitaki KD, Xu YJ, Wang Q, Lin JJ-C, Chen Z-Y, Corey DP. 2015. XIRP2, an actin-binding protein essential for inner ear hair-cell stereocilia. Cell Reports 10:1811–1818. DOI: 10.1016/j.celrep.2015.02.042, PMID: 25772365

Scheibinger M, Janesick A, Benkafadar N, Ellwanger DC, Jan TA, Heller S. 2022. Cell-type identity of the avian utricle. Cell Reports 40:111432. DOI: 10.1016/j.celrep.2022.111432, PMID: 36170825

Schulz-Mirbach T, Ladich F. 2016. Diversity of Inner Ears in Fishes: Possible Contribution Towards Hearing Improvements and Evolutionary Considerations. Advances in Experimental Medicine and Biology 877:341–391. DOI: 10.1007/978-3-319-21059-9_16, PMID: 26515322

Sheets L, Kindt KS, Nicolson T. 2012. Presynaptic CaV1.3 channels regulate synaptic ribbon size and are required for synaptic maintenance in sensory hair cells. The Journal of Neuroscience: The Official Journal of the Society for Neuroscience 32:17273–17286. DOI: 10.1523/JNEUROSCI.3005-12.2012, PMID: 23197719

Shi T, Beaulieu MO, Saunders LM, Fabian P, Trapnell C, Segil N, Crump JG, Raible DW. 2023. Single-cell transcriptomic profiling of the zebrafish inner ear reveals molecularly distinct hair cell and supporting cell subtypes. eLife 12:e82978. DOI: 10.7554/eLife.82978, PMID: 36598134

Shi T, Kim Y, Llamas J, Wang X, Fabian P, Lozito TP, Segil N, Gnedeva K, Crump JG. 2024. Long-range Atoh1 enhancers maintain competency for hair cell regeneration in the inner ear. Proceedings of the National Academy of Sciences of the United States of America 121:e2418098121. DOI: 10.1073/pnas.2418098121, PMID: 39671177

Shin J-B, Krey JF, Hassan A, Metlagel Z, Tauscher AN, Pagana JM, Sherman NE, Jeffery ED, Spinelli KJ, Zhao H, Wilmarth PA, Choi D, David LL, Auer M, Barr-Gillespie PG. 2013. Molecular architecture of the chick vestibular hair bundle. Nature Neuroscience 16:365–374. DOI: 10.1038/nn.3312, PMID: 23334578

Sprague J, Clements D, Conlin T, Edwards P, Frazer K, Schaper K, Segerdell E, Song P, Sprunger B, Westerfield M. 2003. The Zebrafish Information Network (ZFIN): the zebrafish model organism database. Nucleic Acids Research 31:241–243. DOI: 10.1093/nar/gkg027, PMID: 12519991

Stone JS, Cotanche DA. 2007. Hair cell regeneration in the avian auditory epithelium. The International Journal of Developmental Biology 51:633–647. DOI: 10.1387/ijdb.072408js, PMID: 17891722

Stone JS, Leaño SG, Baker LP, Rubel EW. 1996. Hair cell differentiation in chick cochlear epithelium after aminoglycoside toxicity: in vivo and in vitro observations. The Journal of Neuroscience: The Official Journal of the Society for Neuroscience 16:6157–6174. DOI: 10.1523/JNEUROSCI.16-19-06157.1996, PMID: 8815898

Sur A, Wang Y, Capar P, Margolin G, Prochaska MK, Farrell JA. 2023. Single-cell analysis of shared signatures and transcriptional diversity during zebrafish development. Developmental Cell 58:3028–3047.e12. DOI: 10.1016/j.devcel.2023.11.001, PMID: 37995681

Tao L, Yu HV, Llamas J, Trecek T, Wang X, Stojanova Z, Groves AK, Segil N. 2021. Enhancer decommissioning imposes an epigenetic barrier to sensory hair cell regeneration. Developmental Cell 56:2471–2485.e5. DOI: 10.1016/j.devcel.2021.07.003, PMID: 34331868

van der Valk WH, van Beelen ESA, Steinhart MR, Nist-Lund C, Osorio D, de Groot JCMJ, Sun L, van Benthem PPG, Koehler KR, Locher H. 2023. A single-cell level comparison of human inner ear organoids with the human cochlea and vestibular organs. Cell Reports 42:112623. DOI: 10.1016/j.celrep.2023.112623, PMID: 37289589

Wallis D, Hamblen M, Zhou Y, Venken KJT, Schumacher A, Grimes HL, Zoghbi HY, Orkin SH, Bellen HJ. 2003. The zinc finger transcription factor Gfi1, implicated in lymphomagenesis, is required for inner ear hair cell differentiation and survival. Development (Cambridge, England) 130:221–232. DOI: 10.1242/dev.00190, PMID: 12441305

Walls WD, Azaiez H, Smith RJH. n.d. Hereditary Hearing Loss Homepage. https://hereditaryhearingloss.org.

Wan G, Corfas G, Stone JS. 2013. Inner ear supporting cells: rethinking the silent majority. Seminars in Cell & Developmental Biology 24:448–459. DOI: 10.1016/j.semcdb.2013.03.009, PMID: 23545368

Wang T, Ling AH, Billings SE, Hosseini DK, Vaisbuch Y, Kim GS, Atkinson PJ, Sayyid ZN, Aaron KA, Wagh D, Pham N, Scheibinger M, Zhou R, Ishiyama A, Moore LS, Maria PS, Blevins NH, Jackler RK, Alyono JC, Kveton J, Navaratnam D, Heller S, Lopez IA, Grillet N, Jan TA, Cheng AG. 2024a. Single-cell transcriptomic atlas reveals increased regeneration in diseased human inner ear balance organs. Nature Communications 15:4833. DOI: 10.1038/s41467-024-48491-y, PMID: 38844821

Wang T, Yang T, Kedaigle A, Pregernig G, McCarthy R, Holmes B, Wu X, Becker L, Pan N, So K, Chen L, He J, Mahmoudi A, Negi S, Kowalczyk M, Gibson T, Druckenbrod N, Cheng AG, Burns J. 2024b. Precise genetic control of ATOH1 enhances maturation of regenerated hair cells in the mature mouse utricle. Nature Communications 15:9166. DOI: 10.1038/s41467-024-53153-0, PMID: 39448563

Wilkerson BA, Zebroski HL, Finkbeiner CR, Chitsazan AD, Beach KE, Sen N, Zhang RC, Bermingham-McDonogh O. 2021. Novel cell types and developmental lineages revealed by single-cell RNA-seq analysis of the mouse crista ampullaris. eLife 10:e60108. DOI: 10.7554/eLife.60108, PMID: 34003106

Wilmarth PA, Krey JF, Shin J-B, Choi D, David LL, Barr-Gillespie PG. 2015. Hair-bundle proteomes of avian and mammalian inner-ear utricles. Scientific Data 2:150074. DOI: 10.1038/sdata.2015.74, PMID: 26645194

Wu J, Zhang Y, Mao S, Li W, Li G, Li H, Sun S. 2023. Cross-species analysis and comparison of the inner ear between chickens and mice. The Journal of Comparative Neurology 531:1443–1458. DOI: 10.1002/cne.25524, PMID: 37462291

Wu T, Hu E, Xu S, Chen M, Guo P, Dai Z, Feng T, Zhou L, Tang W, Zhan L, Fu X, Liu S, Bo X, Yu G. 2021. clusterProfiler 4.0: A universal enrichment tool for interpreting omics data. Innovation (Cambridge (Mass.)) 2:100141. DOI: 10.1016/j.xinn.2021.100141, PMID: 34557778

Xiang M, Gan L, Li D, Chen ZY, Zhou L, O’Malley BW, Klein W, Nathans J. 1997. Essential role of POU- domain factor Brn-3c in auditory and vestibular hair cell development. Proceedings of the National Academy of Sciences of the United States of America 94:9445–9450. DOI: 10.1073/pnas.94.17.9445, PMID: 9256502

Xu Z, Tavakoli A, Kulasooriya S, Liu H, Tu S, Bloom C, Li Y, Johnson TD, Zuo J, Tao L, Kachar B, He DZ. 2026. The dual molecular identity of vestibular kinocilia bridges structural and functional traits of primary and motile cilia. eLife 14:RP108071. DOI: 10.7554/eLife.108071, PMID: 42029006

Zheng J, Shen W, He DZ, Long KB, Madison LD, Dallos P. 2000. Prestin is the motor protein of cochlear outer hair cells. Nature 405:149–155. DOI: 10.1038/35012009, PMID: 10821263

Zheng L, Sekerková G, Vranich K, Tilney LG, Mugnaini E, Bartles JR. 2000. The deaf jerker mouse has a mutation in the gene encoding the espin actin-bundling proteins of hair cell stereocilia and lacks espins. Cell 102:377–385. DOI: 10.1016/s0092-8674(00)00042-8, PMID: 10975527

